# Diverse conjugative elements silence natural transformation in *Legionella* species

**DOI:** 10.1101/623074

**Authors:** Isabelle Durieux, Christophe Ginevra, Laetitia Attaiech, Kevin Picq, Pierre-Alexandre Juan, Sophie Jarraud, Xavier Charpentier

## Abstract

Natural transformation, *i.e.* the uptake of DNA and its stable integration in the chromosome, is a major mechanism of horizontal gene transfer and is common in bacteria. The vast majority of bacterial genomes carry the specific genes involved in natural transformation, yet only a fraction of species are deemed naturally transformable. This is typically explained by the inability of standard laboratory conditions to induce this phenotypic trait. However, even when the inducing conditions are known, large intraspecific variations have been reported. In this study, we investigated the conservation and distribution of natural transformability in the human pathogen *Legionella pneumophila*. Using a panel of 113 clinical isolates, we found that natural transformability is relatively conserved but shows large variations inconsistent with the phylogeny. By conducting a genome-wide association study (GWAS) we identified the conjugative plasmid pLPL as a source of these intraspecific variations. We further show that the plasmid inhibits transformation by simultaneously silencing the genes required for DNA uptake and recombination, *comEC, comEA, comF* and *comM*. We identified a plasmid-encoded small RNA (sRNA), RocRp, as solely responsible for the silencing of natural transformation. RocRp is homologous to the highly conserved and chromosome-encoded RocR which controls the transient expression of the DNA uptake system. We show that RocRp can take over the function of RocR, by acting as a substitute, ensuring that the bacterial host of the conjugative plasmid does not become naturally transformable. Distinct homologs of this plasmid-encoded sRNA are found in diverse conjugative elements in other *Legionella* species, suggesting that silencing natural transformation is beneficial to these genetic elements. We propose that transformation-interfering factors are frequent genetic cargo of mobile genetic elements, accounting for intraspecific variations in natural transformation but also responsible for the apparent non-transformability of some species.

## Introduction

One of the remarkable feature of bacteria is their ability to naturally undergo genetic transformation. This property is linked to the unique ability of bacteria to import exogenous DNA and integrate it in their chromosome through homologous recombination (1). Despite major difference in their cell wall architecture, the process is generally conserved between Gram-positive and Gram-negative bacteria (2). It first involves an extracellular type IV pilus which is proposed to serve as a ratchet for exogenous DNA (3–5). Once at the cytoplasmic membrane, the captured DNA is converted into a single strand DNA (ssDNA) molecule (6–8) and transported to the cytoplasm where it is brought to the chromosome for recombination. The entire process relies on the tightly regulated and concerted expression of the type IV pilus and of several proteins essential from importing the DNA to promoting its integration in the chromosome, a situation known as competence (9). Collectively forming a DNA uptake system, these proteins include the periplasmic ComEA protein which interacts with double-strand DNA (10, 11) and ComEC, a putative transmembrane channel for ssDNA (12). Crossing the inner membrane requires ComFA, a cytoplasmic ATPase unique to Gram-positive bacteria (13), and ComFC (denoted ComF in Gram-negative bacteria) (14). Both were recently found to from a complex and interact with DprA (15) which protects the incoming ssDNA and promote its RecA-dependent recombination with the chromosome (16, 17). A DNA helicase (RadA in Gram-positive, ComM in Gram-negative) then facilitate recombination over long distance (18, 19). Among all mechanisms of horizontal gene transfer, which are found to be increasingly diverse (20, 21), natural transformation stands out as being the most conserved mechanism. The core components of the DNA uptake system are encoded by the vast majority of bacterial genomes, with for instance, ComEC found in over 95% of bacterial genomes (22, 23). However, only a fraction of species are deemed naturally transformable (24). This is commonly explained by the inability of standard laboratory conditions to induce the concerted expression of the type IV pilus and the DNA uptake system. This expression, *i.e.* the competence state, is generally transient and triggered by signals that appear as species-specific (1, 25). Interestingly, and as illustrated with *Pseudomonas stutzeri*, closely-related species sometime share the ability to undergo natural transformation while others, such as *P. aeruginosa*, do not take up DNA under the same conditions and then become widely accepted as non-transformable (26). This apparent lack of transformability is also found at shorter phylogenetic distance. Even in well-established transformable species such as *Streptococcus pneumoniae, P. stutzeri* and *Haemophilus influenzae*, from 30 to 60% of strains consistently fail to transform (27–31). The underlying cause of this phenotypic heterogeneity is not understood. We here sought to explore this phenomenon in *Legionella pneumophila*, a human pathogen whose genome is shaped by high rates of recombination (32–35). *L. pneumophila* was first found to be naturally transformable under microaerophilic conditions (36). Natural transformation is constitutive in the lab-evolved strain lp02 when grown at 30°C, while its Philadephia-1 parent isolate only showed low transformability levels (37). Characterization of the transformability trait the undomesticated clinical isolate Paris revealed that natural transformability is transient at 30°C, occurring at the transition of the exponential growth phase and the stationary phase (38, 39). The high rates of recombination inferred from genomic analysis would be consistent with the trait of natural transformability. Yet, all strains reported as transformable belong to the same serogroup and the degree of conservation of the transformability trait has not been investigated.

We here document the distribution and variability of natural transformability in a panel of clinical isolates of *L. pneumophila* belonging to 42 sequence type (ST). While the trait is displayed by the majority of isolates, examination of the non-transformable isolates revealed the pervasive inhibition of natural transformation by a single conjugative element. Several other conjugative elements have adopted a similar strategy tailored to *Legionella* species. We here establish that sparsely-distributed mobile genetic elements conflicting with natural transformation can contribute to instraspecific variations in natural transformability. We propose that a widespread distribution of such elements may also cause the apparent lack of transformability of specific species.

## Results

### Large instraspecific variations in natural transformability are incongruent with the phylogeny

We initially determined the natural transformability of a random set of 25 clinical isolates, along with the Paris and Lens isolates, all belonging to 12 different STs (Fig. 1A). As expected, the Paris strain and closely-related ST1 isolates showed high transformability levels (average transformation frequencies ranging from 10^−6^ to 10^−5^). Interestingly, some distantly-related strains, such as LG-0724-2038 (ST23) also display high transformability levels. Out of the 27 tested isolates, 20 showed transformation frequencies that were consistently greater than the detection limit (10^−9^), indicating that natural transformability is a conserved trait in *L. pneumophila*. In contrast, some strains were poorly transformable or even consistently unable to generate transformants, as for instance the Lens isolate. Importantly, and as exemplified by isolates HL-0647-5014 and HL-0640-5008, these strong variations in natural transformability, spanning four order of magnitude, are incongruent with the phylogeny. In order to gain insight into these extensive variations we tested natural transformability of an extended set of 109 isolates (40), including the same 25 clinical isolates, Paris and Lens isolates, the Olda and 130b strains (Table S1). For this extended set, now encompassing 42 STs, transformability was scored from 0 to 3 using a higher-throughput assay (see Material and Methods) and averaged from four independent experiments (Table S1). Phylogenetic analysis using core-genome polymorphism confirmed that the phenotype of natural transformability has mixed relationships with phylogeny (Fig. 1B). For instance, all isolates belonging to the branch of the Paris strain are consistently transformable. This is concordant with the high rates or recombination observed in this ST1 lineage, with an average of nearly 10% of the genome of isolates originating from recombination (34). Similarly, all isolates of the ST47 cluster are consistently poorly transformable. Again, this is consistent with the clonal nature of the ST47 lineage, showing no signs of recombination events (34). In contrast, in other branches, non-transformable isolates are mixed with transformable isolates. This is particularly noticeable in the cluster of strains belonging to the ST23 group which incudes an equal number of non-transformable and highly transformable isolates. Overall, natural transformability is a conserved trait. Yet, the natural transformation phenotype shows extensive variations that are not strictly associated with phylogenetic relationships. This phenomenon is consistent with observations in other transformable species for which the underlying cause have remained elusive (27–30, 41).

**Figure 1.**
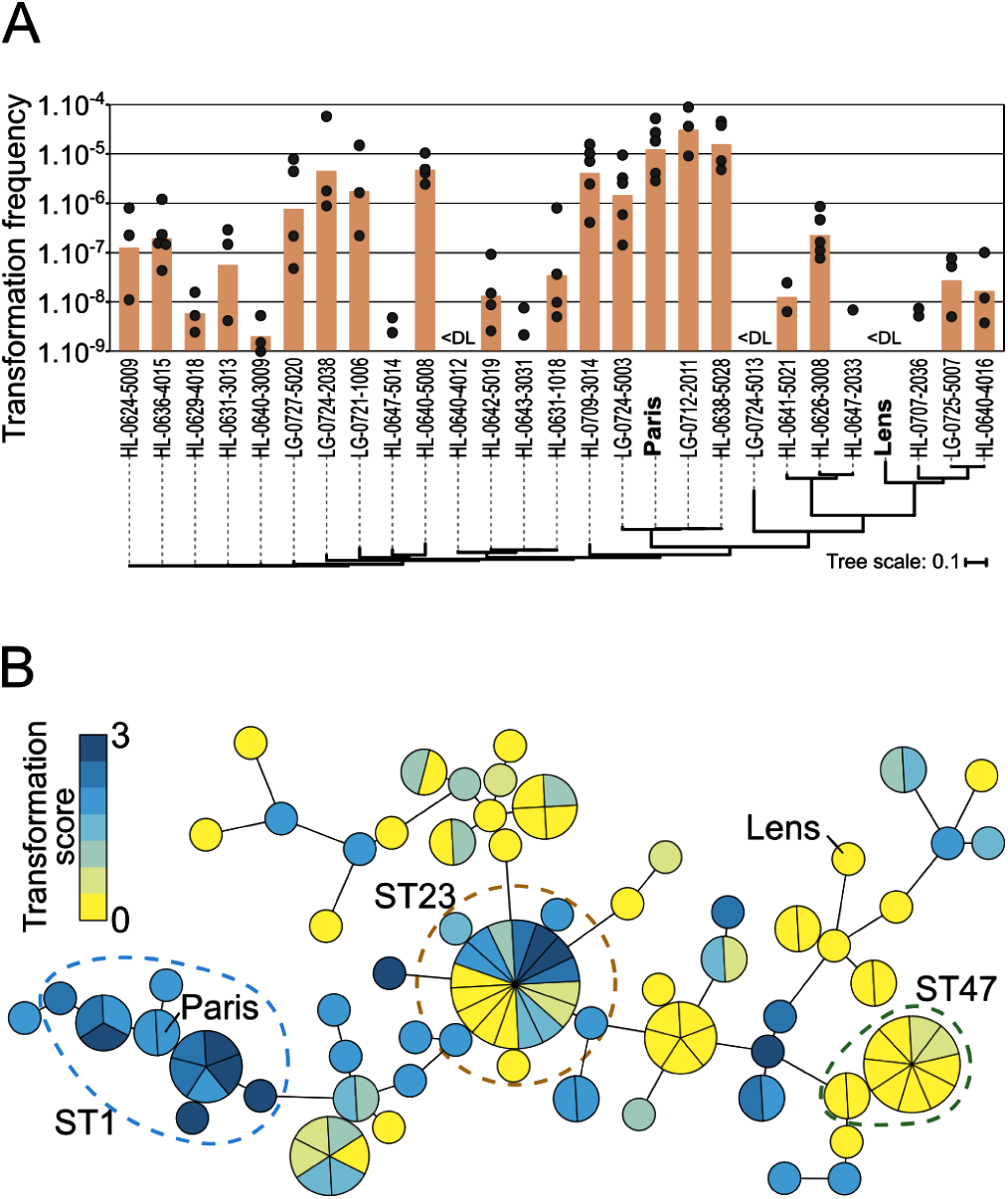
Extensive variations of the trait of natural transformation are inconsistent with the phylogeny. A. Quantitative analysis of natural transformability of 25 isolates of *L. pneumophila*. Natural transformation was tested by incubating each isolate with a non-replicative plasmid carrying the *ihfB* gene interrupted by a gene conferring resistance to kanamycin. Each isolate was tested from three to five times on independent occasions. Bars represent the geometric mean when at least two frequencies (black dots) could be determined. <DL, below detection limit. A region homologous to the transforming DNA is present in all tested isolates. Phylogenetic relationships were computed based on core-genome SNPs and displayed as a phylogram. B. Semi-quantitative analysis of natural transformability of 113 isolates of *L. pneumophila*. Transformability was determined in 96-well format using as transforming DNA a 4 kb PCR product encompassing a *rpsL* allele conferring resistance to streptomycin. Following incubation with DNA for at least 24H, 10 µL of the cultures were spotted on plates containing streptomycin. Transformation was scored from 0 to 3 as a function of the approximate number of colonies that developed in the spot (see Materials and Methods). Transformation scores were determined four times independently. A control without DNA was conducted in parallel. For each experiment, a score was retained only if it was superior to the score determined in the no DNA conditions. The median score is reported for the *n* number of determinations that met this criteria (See Table S1). Genetic relationship were determined by cgMLST and visualized using a minimum spanning tree displaying transformation scores color coded from 0 (yellow) to 3 (dark blue).

### GWAS associates the sparsely distributed conjugative plasmid pLPL with non-transformability

We hypothesize that genetic factors were responsible for the extensive variations of natural transformability. The panel of 113 isolates offered the possibility to reveal such factor through the statistical associations between genetic polymorphism and the transformation phenotype. We thus converted the transformation scores into a binary phenotype (transformable T, score 1-3 or non-transformable NT, score 0-0.5) (Fig. S1A) and conducted a genome-wide association studies (GWAS) using DBGWAS (42). DBGWAS provides a graphical output that can visually distinguish genetic determinants associating with phenotypes that correspond to single-nucleotide polymorphism (SNP) or to horizontal gene transfer (HGT) events. The output of DBGWAS on our dataset is typical of an event of HGT with the top three subgraphs (q-value<0.1) displaying nearly linear structures consisting of respectively 683, 51 and 93 unambiguous sequences that individually associate with the NT phenotype. The combined 827 sequences mapped exclusively onto the sequence of pLPL (Fig. S1A), a plasmid that was originally identified in the Lens strain (43). Plasmid pLPL is a large element (59.8 kb) which notably carries a conjugative system and a Type I-F CRISPR-cas immunity system also found in the chromosome of this strain (Fig. S1A). The pLPL sequence is found in one third of the non-transformable isolates with 14 isolates carrying a pLPL variant (Fig. S1B). Of these, 10 are essentially identical to the original pLPL plasmid of the Lens isolate (Fig. S2). Other plasmids show large insertions and deletions, including the region carrying the CRISPR-Cas locus. Analysis of 537 available *L. pneumophila* genomes revealed that 5.4% of isolates carry a pLPL element (Fig. 2A). As a mean of comparison, we also searched for the presence of pLPP, a distinct plasmid initially described in the Paris isolate (43). Plasmid pLPP was found in 6.9% of isolates and seems restricted to clinical isolates belonging to the ST1 group. In contrast, pLPL elements are sparsely distributed throughout the *L. pneumophila* isolates, suggesting that they are horizontally spreading by conjugation. We thus tested the functionality of the conjugative system of the pLPL plasmid found in isolate HL-0640-3009, hereafter denoted pLPL^3009^ and strain 3009, respectively. In strain 3009, we generated pLPL^3009KF^ which carries the selectable/counter-selectable “kan-mazF” cassette (see Material and Methods) in a pseudogene (corresponding to *plpp0008* in the Lens pLPL). In co-culture experiments with the non-transformable laboratory strain JR32, we found that pLPL^3009KF^ could transfer at a frequency of 3.2±1.18×10^−6^ (transconjugants per recipient). We could also transfer pLPL^3009KF^ by conjugation to the Paris strain and two other isolates, LG-0712-2011 and HL-0638-5028, that display high transformability levels (Fig. 1A). In all three strains, acquisition of pLPL^3009KF^ decreased transformation frequencies by three order of magnitude, below the detection limit (Fig. 2B). The counter-selectable kan-mazF cassette of pLPL^3009KF^ allowed us to isolates clones of strain 3009 that spontaneously lost it. Clones lacking pLPL occur at a frequency of ~10^−5^ and plasmid extraction confirmed the absence of pLPL in strain 3009ΔpLPL (Fig. S3). Cured of pLPL^3009KF^, the strain can now efficiently undergo natural transformation (Fig. 2C). These results experimentally confirm the GWAS result that pLPL is a potent inhibitor of natural transformation.

**Figure 2.**
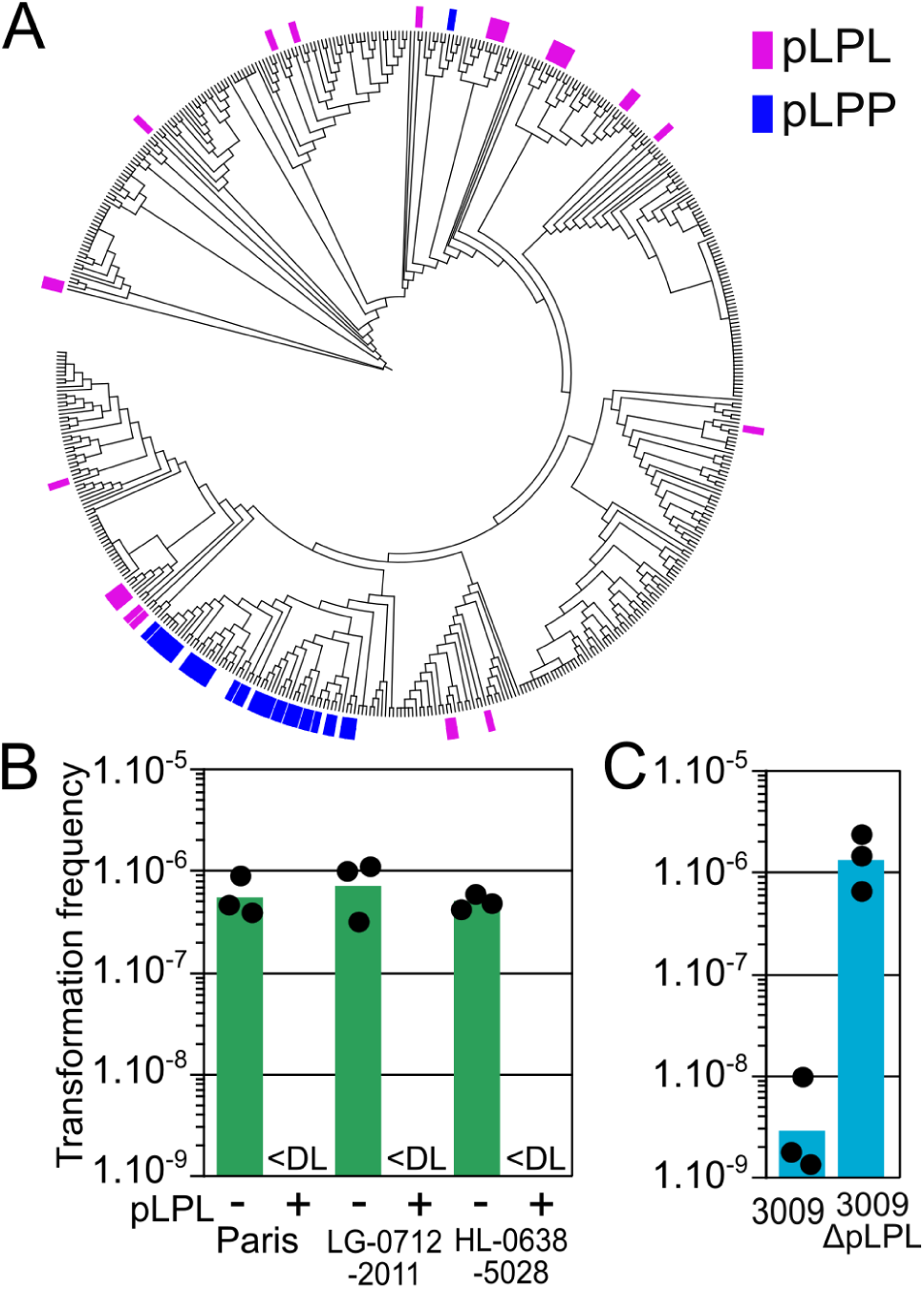
Plasmid pLPL is a sparsely distributed inhibitor of natural transformation. A. Cladogram based on core-genome SNPs of 537 *L. pneumophila* genomes available from public repositories. Presence or absence of pLPL (blue) and pLPL (pink) was determined using Seqfindr and displayed on the cladogram drawn with iTol. B. Natural transformability of Paris and clinical isolates compared to their transconjugants carrying pLPL from isolate 3009. Streptomycin-resistant mutants of the tested strains were first isolated and used as recipients in conjugation experiments with strain 3009 carrying pLPL^3009KF^. Natural transformation was tested by incubation with genomic DNA from a Paris strain carrying a chromosomal gentamycin resistance gene (Paris_H1). Data points are from three independently performed experiments and bars represent the geometric mean of the transformation frequencies. <DL, below detection limit. C. Natural transformability of strain 3009 and 3009ΔpLPL which lack pLPL. Transformation was tested by incubation with a PCR product of the *rpsL* allele conferring resistance to streptomycin, similarly to as in Figure 1A. Data points are from three independently performed experiments and bars represent the geometric mean of the transformation frequencies.

### Plasmid pLPL inhibits natural transformation by silencing the DNA uptake system

Natural transformability is a transient phenotype in *L. pneumophila*, occurring at the onset of the stationary phase. Transformability correlates with expression of the genes encoding the DNA uptake system, notably *comEA* whose expression is a marker of natural transformability (38, 44, 45). In the Paris strain, expression of *comEA* is repressed in exponential phase, expressed at the transition between exponential to stationary phase and then repressed again in stationary phase (38). Northern-blot analyses of *comEA* expression in the original isolates 3009 shows that *comEA* is never expressed during growth (Fig. 3A). However, curing pLPL restored a transient expression pattern of *comEA*. Thus, pLPL appears to inhibit expression of *comEA*. RNAseq transcriptional profiling of the 3009 and 3009ΔpLPL strains shows that pLPL^3009^ also prevents expression of *comF, comEC, comM* and *radC* (Fig. 3B, *x* axis and Table S2). The concerted targeted inhibition of these five genes is reminiscent of the action of RocR, a *Legionella*-specific but highly conserved 66 nt-long small non-coding RNA (sRNA) that silences these genes in exponential phase (38). RocR depends on the RocC RNA chaperone to form a duplex with the 5’ untranslated region of the genes encoding the DNA uptake system, preventing translation and exposing the targeted mRNA to cellular nucleases. Absence of RocR or RocC result in a constitutively transformable phenotype (38). A blast search using the RocR sequence revealed a sequence with 80% identity and located in an intergenic region of pLPL plasmids (red bar between 35 and 40 kb, Fig. S1). This sequence is missing in the pLPL variants of pLPL-positive isolates HL-0637-4025 and HL-0703-5020 that appeared transformable (Fig. S2). Rapid amplification of cDNA ends (RACE) confirmed transcription initiating at three positions, generating three variants of 65, 68 and 70 nt (Fig. S4). We termed this gene *rocRp* for “*rocR <u>p</u>*lasmidic”. RNAseq transcriptional profiling of a deletion mutant 3009Δ*rocRp* shows overexpression of the same genes induced in the 3009ΔpLPL strain (Fig. 3B and Table S2). Thus, *rocRp* alone is responsible for the repression of the DNA uptake system caused by pLPL. Accordingly, strain 3009Δ*rocRp* can undergo natural transformation, similarly to strain 3009ΔpLPL (Fig. 3C). We then inserted the *rocRp* gene in the plasmid pLPP, naturally carried by the Paris strain. The resulting strain Paris pLPP::*rocRp* is no longer transformable, demonstrating that *rocRp* is sufficient to inhibit natural transformation (Fig. 3D). In addition, *rocRp* can also fully repress the constitutive and elevated transformability of a Δ*rocR* mutant, indicating that RocR and RocRp fulfill the same function. Consistent with this result, and just like RocR, the RocRp RNA appears to require the RNA chaperone RocC as it cannot repress natural transformation of the *rocC*TAA mutant (Fig. 3D). In conclusion, pLPL inhibits transformation by encoding an sRNA that mimics the chromosome-encoded sRNA RocR and silence the genes encoding the DNA uptake system.

**Figure 3.**
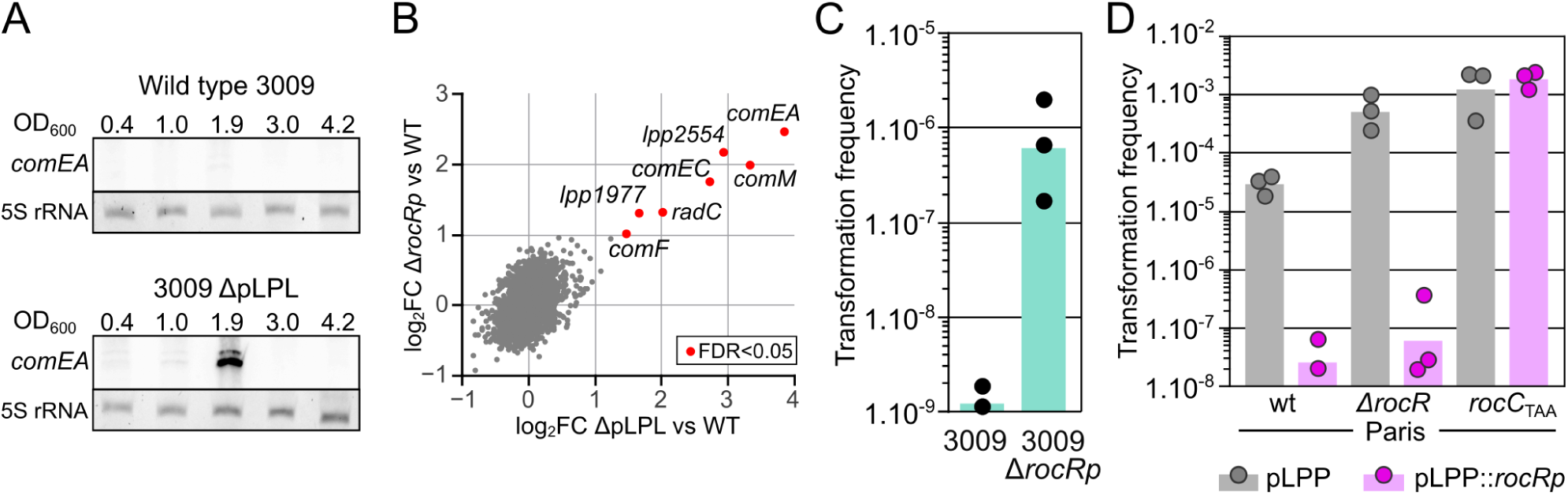
Plasmid pLPL silences the DNA uptake system by expressing the sRNA RocRp. A. Northern-blot analysis of *comEA* expression during growth of strains 3009 and 3009ΔpLPL in AYE medium at 30°C. Total RNA were extracted at the indicated optical densities (measured at 600 nm, OD_600_) of the culture, and corresponding to the exponential growth phase (0.4 to 1.0), the transition phase (1.9) and early (3.0) and late stationary phase (4.2). Total RNA was separated on an 8% denaturing polyacrylamide gel and *comEA* transcripts were revealed with a biotinylated oligonucleotide probe. 5S rRNA was used as a loading control. B. RNAseq transcriptional profiling of strains 3009ΔpLPL and 3009Δ*rocRp* compared to wild-type 3009. Total RNA was extracted from cultures collected at an optical density of 1.8-2.0 (at which expression of *comEA* was observed). Following rRNA depletion, RNAs were reverse-transcribed and sequenced (Illumina). Each set of strains (3009ΔpLPL, 3009 Δ*rocRp* and wt) was grown in three independent experiments (n=3). The number of reads was normalized and enrichment is reported as log _2_-transformed fold-change (log_2_FC). Benjamin-Hochberg correction was applied to P-value (false-discovery rate, FDR). Individual genes (gray dots) were considered deferentially expressed if log_2_FC was >1 or <-1 and if FDR<0.05 (red dots). C. Natural transformability of strain 3009 and 3009Δ*rocRp*. Transformation was tested by incubation with a PCR product of the *rpsL* allele conferring resistance to streptomycin, similarly to as in Figure 1A. Data points are from three independently performed experiments and bars represent the geometric mean of the transformation frequencies. D. Natural transformability of the Paris strain and isogenic mutants for *rocC* (*rocC*_*TAA*_) and RocR (Δ*rocR*) in which the *rocRp* gene was introduced in the pLPP plasmid (pink) or not (grey). Transformation was tested by incubation with a PCR product of the *rpsL* allele conferring resistance to streptomycin, similarly to as in panel C. Data points are from three independently performed experiments and bars represent the geometric mean of the transformation frequencies.

### RocRp acts as a substitute for RocR

How does RocRp exert a dominant effect over the chromosome-encoded RocR? We investigated this situation by looking at the expression of one of their targets, *comEA*. We previously found that during the transition phase, the decreased expression level of RocR relieves silencing, allowing for the transient expression of *comEA* (Fig. 4A and (38)). However, in the strain Paris pLPP::*rocRp, comEA* is silenced in the transition phase suggesting that RocRp may be expressed differently than RocR. Indeed, expression of RocRp from the pLPP plasmid shows a pattern in negative of that of RocR (Fig. 4A). Expression of RocRp is low in exponential phase but increases at the end of the exponential phase to reach its maximal level at the transition phase (Fig. 4A). This expression pattern of RocRp is also observed when it is expressed from its original pLPL plasmid in strain 3009, despite the fact that strain 3009 reaches stationary phase at a lower OD (~4) than the Paris strain (~6) (Fig. S5A). Both RocR and RocRp are undetectable in late stationary phase (Fig. 4A), possibility because of the absence of RocC in stationary phase (OD>4.5) (Fig. S5B), which is needed to stabilize RocR (38). The northern-blot analysis indicates that RocRp levels in the transition phase compensate for the decreased levels of RocR. Yet, northern-blot analysis are not suited to determine the relative levels of RocR and RocRp. To determine the relative levels of RocR and RocRp engaged in silencing of the DNA uptake system, we immunopurified RocC from cultures at OD 0.9-1 and OD 2.5-3. RNA bound to RocC were purified and the relative abundance of RocR and RocRp was determined using a restriction analysis following a reverse-transcription step (See Material and Methods). In exponential phase (OD 0.9-1), RocC binds three times more RocR than RocRp but in the transition phase this ratio shifts in favor of RocRp which becomes the most abundant sRNA bound to RocC (Fig. 4B). Interestingly, the half-life of RocR and RocRp correlates with their relative association with RocC. In exponential phase, when RocC mainly binds RocR, RocR shows a long half-life of 330±22 minutes (n=3) which decreases by 3-fold to 109±65 minutes in the transition phase. RocRp seems less stable than RocR, but shows the opposite pattern with a half-life of 36±7 minutes in exponential phase that increases 3-fold to 107±39 minutes when it is bound to RocC in the transition phase (Fig. S5C). The results are consistent with our previous finding that RocC stabilizes the ncRNA (38) and suggest that the engagement of RocRp in RocC-assisted silencing increases its stability. To test this hypothesis we analyzed the expression and stability of RocRp in a Δ*rocR* mutant. RocRp can effectively silence natural transformation in the absence of RocR (Fig. 3D). In the absence of RocR, RocRp remains expressed in the transition phase but now shows high expression levels in exponential phase, behaving like RocR (Fig. 3C). Indeed, in exponential phase RocRp now shows a half-life of 412±91 minutes, similar to that of RocR (Fig. S5D). This indicates that the stability of RocRp is not only dictated by its intrinsic properties but rather by its involvement in RocC-dependent silencing. If RocRp levels are affected by the absence of RocR, in contrast, the presence of RocRp does not significantly alter the expression levels of RocR, indicating that RocRp cannot displace RocR from RocC (Fig. 4A). Altogether the data suggest that RocRp does not compete with RocR. Rather, RocRp fulfills the same function as RocR, but only when the latter is missing. Thus, RocRp appears to act as a substitute, taking over the function of RocR if its levels were to decrease.

**Figure 4.**
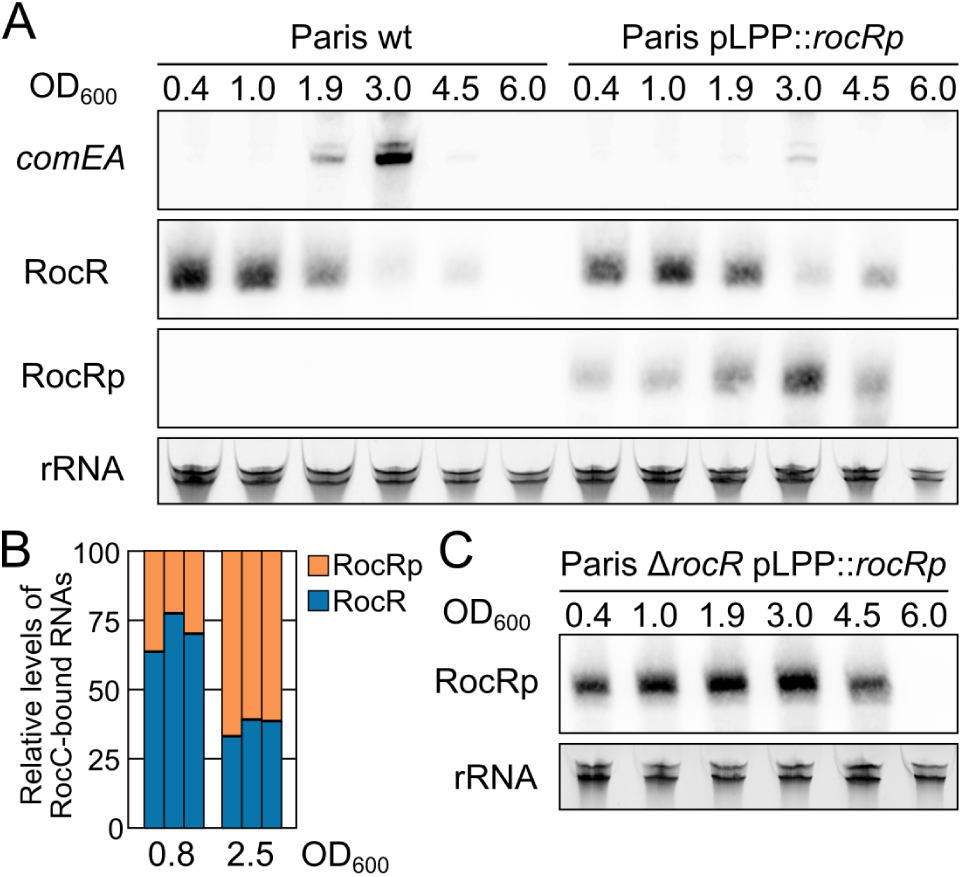
RocRp acts as a substitute for RocR. A. Northern-blot analysis of *comEA* mRNA, RocR and RocRp expression during growth of the wild-type Paris strain or carrying the *rocRp* gene in pLPP (Paris pLPP::*rocRp*) in AYE medium at 30°C. Total RNA was extracted at the indicated optical densities (OD_600_) during the exponential growth phase (0.4 to 1.0), the transition phase (1.9 and 3.0) and the early (4.5) and late stationary phases (6.0). Ribosomal RNA (rRNA) were used as loading control. B. Relative quantification of RocR and RocRp bound to RocC in exponential phase (OD=0.8) and transition phase (OD=2.5). For each OD, three independent cultures of the strain Paris pLPP::*rocRp* were obtained. Bacteria were lysed and RocC was purified with RocC-specific antibodies. RNA that co-purified with RocC were reverse transcribed with a mix of primers specific for RocR and RocRp, each bearing the same 5’ end extension. The resulting cDNA were amplified by PCR using primers corresponding to the 5’ extension of the RT primer and a reverse primer that matches both RocR and RocRp. Amplification of either RocR or RocRp produces a 85 bp DNA. PCR products originating from RocR and RocRp were distinguished by restriction with NruI, which cuts only the RocRp PCR product in two fragments. The restriction products were analyzed by capillary electrophoresis, allowing for the relative quantification of fragments corresponding to RocR and RocRp. C. Northern-blot analysis of RocRp expression during growth in AYE medium at 30°C of the strain Paris pLPP::*rocRp* deleted of *rocR*. Ribosomal RNAs (rRNA) were used as loading controls.

### A diverse set of conjugative elements in the *Legionella* genus carry RocRp homologs

RocR is highly conserved in the *Legionella* genus. RocR was found in 833 out of 835 *Legionella*/*Fluoribacter*/*Tatlockia* genome assemblies. One assembly in which RocR is missing is incomplete, the other one belong to a lice symbiont with drastically reduced genome size (1/5 of a typical *Legionella* genome). In addition to being highly conserved, its sequence is uniquely invariant, showing no SNP in all *L. pneumophila* isolates and in 97 out 138 non-*pneumophila* isolates. Only 38 non-*pneumophila* species isolates showed one SNP and three showed two SNPs. The ubiquity of RocR in *Legionella* species suggests that in all species natural transformation is regulated by this non-coding RNA. Consequently, the inhibition strategy used by pLPL may also function in non-*pneumophila* species. Thus, other mobile genetic elements could also use a RocRp homologs to interfere with natural transformation. Blast search on publicly available genomes revealed three distinct RocRp homologs in an unclassified *Legionella* species (km542) and in *L. geestiana* (DSM21217) and in *L. israelensis* (DSM19235) type strains (Fig. 5A). The RocRp homologs of *L. sp.* km542 and *L. israelensis* are closely related to the RocRp from pLPL, whereas the RocRp from *L. geestiana* is more closely related to the chromosomal RocR (Fig. 5B). All RocRp homologs were found to be encoded by intergenic regions (Fig. 5C). Both the *L. sp.* km542 and *L. geestiana* RocRp homologs are carried by episomal conjugative elements (of 66.4 and 60.9 kb, respectively) while the *L. israelensis* RocRp homolog is part of a 46.6 kb conjugative element integrated in chromosome at the tmRNA-encoding gene. Like pLPL, the plasmid of *L. sp.* km542 encodes a type F conjugative system. In contrast the integrated conjugative element (ICE) of *L. geestiana* DSM21217 and the plasmid of *L. israelensis* DSM19235 encode type T systems (46). All four elements share no other homology (Fig. 5B). We were intrigued by the occurrence of RocRp homologs in *L. geestiana* and in *L. israelensis* for which a single genome is available and were isolated in England and Israel, respectively. These species have not been subsequently isolated. We nonetheless could gather one more *L. geestiana* isolate from France and two isolates of *L. israelensis* from France and Norway. The genomes of all three new isolates were sequenced, revealing that they also displayed RocRp homologs but on distinct mobile genetic elements. In *L. geestiana* HL-0438-2026, the RocRp homolog is found again on a 53.3 kb episomal element but this time encoding a type F conjugative system (Fig. 5B) (46). In *L. israelensis* 09060433301 isolated in Norway, the RocRp homolog belong to an ICE with an unclassified conjugative system previously reported in the *L. pneumophila* genomic islands (LGI) LpcGI-2 and LppGI-2 of *L. pneumophila* (47). In *L. israelensis* HL-0427-4011, the RocR homolog is found in a another LGI inserting through site-specific integration at tRNA genes (47). Except for TraC, this element shows no known conjugative system but encodes a TraK homolog, a protein known to interact with the oriT-DNA (48) which was identified directly usptream of the *traK* gene. Thus, although devoid of recognizable conjugative system, this LGI may still be mobilized by other conjugative systems of plasmids or ICE. Interestingly, the RocRp homologs and the proximal sequences were identical between the mobile genetic elements found in each species, suggesting that they arose by intra-specific transfers. In all cases, RocRp homologs were found in conjugative or mobilizable elements either found in their episomal form or integrated in the chromosome. In conclusion, the strategy to inhibit transformation by encoding a RocRp homolog has been adopted by a diverse set of conjugative and mobilisable elements in *Legionella* species. Importantly, all known isolates of the two species *L. geestiana* and *L. israelensis* are infected with distinct conjugative elements carrying RocRp homologs.

**Figure 5.**
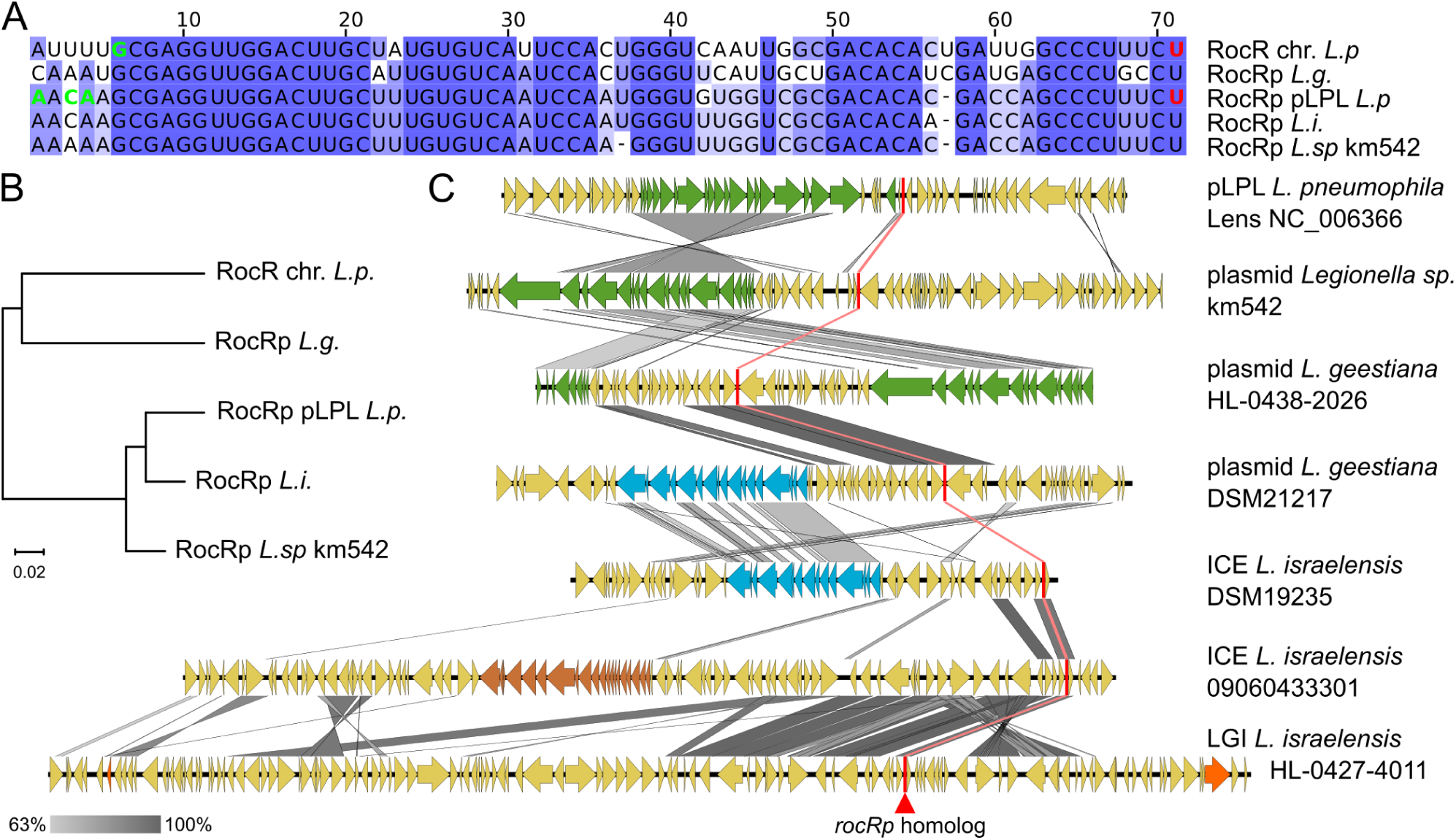
Diverse conjugative elements carry RocRp homologs. A. Sequence alignment of RocR and RocRp homologs found in *Legionella* species. When available, experimentally determined transcription start and termination sites are highlighted with bold green and red nucleotides, respectively. B. Parsimony-based unrooted phylogenetic tree of the genes encoding RocRp homologs. C. Schematic representation of mobile genetic elements carrying a RocRp homolog-encoding gene. Conjugative systems of the type F (similar to the system of F plasmid) and type T (similar to the system of plasmid RP4 or R388) are colored in green and blue, respectively. Incomplete and unclassified conjugative systems are colored in orange and brown, respectively. RocRp homologs are represented as a red bar. Pairwise nucleotide sequence identity is represented by a gradient from light (63%) to dark grey (100%).

## Discussion

We here present the first analysis of the distribution and conservation of natural transformation in *L. pneumophia*. The transformability phenotype is generally conserved, with more 70% of isolates showing transformation frequencies above the detection limit (1.10^−9^) when determined with a resistance marker conferring kanamycin resistance (Fig. 1A). About 45% of isolates (12 out 27) show average transformation frequencies greater than 1.10^−7^. This is comparable to the transformability rate of 52 % in the panel of 113 isolates for which transformability was determined using a less sensitive assay (detection limit >10^−8^) (Fig. 2A). Although conserved, the phenotype is not homogeneously distributed. The ST1 clade groups a majority of transformable isolates, consistent with high recombination rates observed in this clade (33, 34). This contrasts with the absence of transformable isolates in the ST47 clade in which no recombination events could be detected (34). By no means a demonstration, this correlation between transformation and recombination rates yet suggests that natural transformation is a major contributor to genome recombination in *L. pneumophila*. The occurrence of transformable and non-transformable phenotypes in the ST23 clade provided the strongest signal for GWAS and allowed for the identification of pLPL as an inhibitor of natural transformation. We had previously noticed the presence of a sequence with homology to the *rocR* repressor on pLPL, but GWAS provided the first evidence that this sequence was active and could take over the function of RocR. Plasmid pLPL is found in one third of non-transformable isolates and GWAS could not identify other genetic determinants associating with the remaining non-transformable isolates. This may be because different polymorphisms (or diverse mobile genetic elements) can result in the loss of transformability. GWAS also could not identify the underlying reason for the lack of transformability of isolates of the ST47 cluster, which do not carry pLPL. In this instance, the ST47 cluster is homogeneously non-transformable and shows too little genetic polymorphism for a successful GWAS approach.

Tt is striking that a unique conjugative element could be responsible for the lack of transformability of one third of the non-transformable isolates of our panel of 113 isolates. Analysis on a more global scale of over 500 genomes, indicates that at least 5% of *L. pneumophila* isolates are not transformable due to this single conjugative element. This provides the first evidence that conjugative elements significantly contribute to intraspecific variation in natural transformability, a situation observed in other species but so far unexplained (27–30). This is likely not to be unique to *Legionella*, as other instance of transformation inhibition by plasmids and conjugative elements have been sporadically reported in other species. For instance, the high transformability of the *Bacillus subtilis* strains 168, PY79, and JH642 used in laboratories is linked to the loss of plasmid pBS32 which inhibits transformation in their parental strain 3610 (49). This 84 kb plasmid encodes a 30-amino-acid protein ComI needed to inhibit transformation. The exact mechanism by which ComI limits transformation is not known but it is suggested that ComI interferes with the activity the DNA uptake system without altering its expression (49). In contrast, still in *B. subtilis*, the native conjugative plasmid pLS20 does interfere with expression of the DNA uptake system by encoding a homolog of the Rok repressor of ComK, the master transcriptional activator of competence for natural transformation in this species (50). In *Vibrio cholerae*, isolates from the Haiti outbreak are poorly transformable because the VchInd5 ICE encodes a secreted DNAse, IdeA, that degrades exogenous DNA (51). The same VchInd5 ICE with the *ideA* gene was also found in isolates from a site in Bangladesh. It was proposed that the inhibition of natural transformation by a secreted DNAse was co-incidental, resulting from the serendipitous acquisition of a “stowaway” gene (51). The silencing of the DNA uptake system by diverse conjugative elements in *Legionella* appears less fortuitous. First, this interference seems tailored to *Legionella* species, as it relies on a homolog of an sRNA that constitute the core regulator of natural transformation in *Legionella*. Second, it is recurrent, with a handful of conjugative elements encoding distinct RocRp homologs. Taken together, all these active interference now designate natural transformation as a common target of conjugative elements but also of other mobile genetic elements (MGEs). Indeed, a repressor of the ComRS system, which controls competence in *Streptococcus pyogenes*, was recently identified on a prophage of this species (52). Also, several ICE and prophages have been reported to insert and disrupt genes involved in natural transformation ((53, 54) and references therein). This observation came in support of the proposal that natural transformation is a mechanism to cure the genome of parasitic MGEs by homologous recombination with chromosomal DNA released by uninfected siblings (53). Inactivating transformation, either through insertional inactivation or by carrying a transformation-interfering genetic cargo, would prevent MGEs removal and thus favor their persistence in the bacterial community. However, the “anti-curing hypothesis” hardly explains the inhibition of transformation by non-integrative plasmids (such as pLPL, pBS32 and pLS20). Blocking transformation could prevent import of large competing plasmids, yet this process is poorly efficient and plasmids can spread at high rates by conjugation. However, blocking DNA uptake could limit recombination and rearrangements that could jeopardize the plasmid stability or impair its propagation. Further work will be required to test this hypothesis.

Even if the evolutionary benefit of transformation inhibition is elusive, it nonetheless indicates that natural transformation is antagonistic to MGEs. In this respect, inhibition of transformation by MGEs is reminiscent of anti-defense mechanisms of phages (anti-restriction systems, anti-CRISPRs)(55). Co-evolution of conflicting entities result in an evolutionary arms race and diverse strategies of defense and counter-defense, often tailored to a specific pair of conflicting entities (56). From only a handful of cases, it is already apparent that MGEs have evolved diverse mechanisms to interfere with natural transformation, targeting several stages of the process. We speculate that a plethora of transformation inhibiting factors (TIF), frequent genetic cargo of MGEs, are yet to be discovered. These largely evade detection because of their diversity, obscuring the widespread occurrence of natural transformability in bacteria. Conversely, once identified, they may represent evidence of natural transformation activities. This is exemplified here by *L. geestiana* and *L. israelensis*, in which the presence of TIF cargo on distinct conjugative elements is a strong indication of the natural transformability of these species. Also, characterizing new TIF could shed light on the regulation and molecular mechanisms of DNA uptake and recombination. Revealing the intricacies of the conflict between MGEs and natural transformation thus represents a unique opportunity to gain insight into a mechanism of HGT, which has captivated scientists for nearly a century (57).

## Material and Methods

### Bacterial Strains and Growth Conditions

All *L. pneumophila* strains and clinical isolates (see Table S1 and S3) were grown in liquid medium ACES [N-(2-acetamido)-2-aminoethanesulfonic acid]-buffered yeast extract (AYE) or on solid media ACES-buffered charcoal yeast extract (CYE) plates at 30 °C or 37 °C. Liquid cultures were performed in a 13-mL tube containing 3 mL of medium in a shaking incubator (Minitron, Infors HT) at 200 rpm. Alternatively, isolates were grown in 100 µL AYE in 96-well plates sealed with a oxygen-permeable membrane (Sigma-Aldrich). When appropriate, chloramphenicol (Cm), gentamicin (Gent) and IPTG (isopropyl β-D-1-thiogalactopyranoside) were used at 5 µg/mL, 10 µg/mL and 500 µM, respectively. Kanamycin (Kan) was used at 15 µg/mL and streptomyci at 50 µg/mL. Oligonucleotides and plasmids are listed in Table S4.

### DNA manipulation

All PCR for assembling transforming DNA were obtained by proof-reading polymerase (PrimeStar Max, Takara). Genomic DNA from *Legionella* was isolated using the Wizard Genomic DNA Purification Kit or the Maxwell DNA extraction kit (Promega). Plasmids from *L. pneumophila* were isolated from 25 mL of culture using the alkaline lysis method, followed by extraction with phenol:chloroform:isoamyl alcohol (25:24:1) followed by ethanol precipitation. The pellet was treated with RNAse A, ethanol-precipitated and resuspended in 50 µL of Tris-HCl 10 mM, pH 8.

### Strain construction

Full sequence of all genetic constructs are available upon request. (A) Construction of derivatives of HL-0640-3009. Strain 3009 pLPL^3009KF^ was obtained by introducing the kan-mazF cassette (58) at position 9074 (between CDS55 and CDS56) of the pLPL plasmid carried by HL-0640-3009. Isolate HL-0640-3009 was naturally transformed with a PCR product consisting of the kan-mazF cassette flanked by 2 kb of sequences corresponding to the insertion site. Three PCR fragments were generated independently, a 2 kb PCR fragment upstream of the targeted insertion site (primers mazF-pLPL_P1F and mazF-pLPL_P2F), a PCR product of the kan-mazF cassette (primers MazFk7-F and MazFk7-R) and a PCR product of a 2kb fragment downstream of the targeted insertion site (primers mazF-pLPL_P1F and mazF-pLPL_P2F). The PCR products were purified, mixed and the assembled PCR product was amplified with primers mazF-pLPL_P1F and mazF-pLPL_P4F. HL-0640-3009 was then naturally transformed with this DNA construct. Due to the low transformability of the isolate (transformation frequencies ~1E-8), only a handful of transformants were obtained. The transformants are resistant to kanamycin and sensitive to IPTG (because it induces expression of the MazF toxin). Correct insertion was validated by PCR. Strain 3009ΔpLPL was isolated by plating culture of strain 3009 pLPL^3009KF^ on CYE plates containing IPTG. Clones resistant to IPTG and sensitive to kanamycin were found to lack the entire pLPL. Plasmid extraction and PCR analysis confirmed the absence of pLPL. Strain 3009Δ*rocRp* was obtained in two steps. First, the kan-mazF cassette was inserted in the *rocRp* gene by following the same method used for inserting the cassette in between CDS55 and 56 (see above). As described above, a PCR product was assembled with a 2 kb fragment upstream of *rocRp* (PCR with primers DeltaRocRlike_P1 and DeltaRocRlike_P2), a PCR product of the kan-mazF cassette (primers MazFk7-F and MazFk7-R) and a PCR product of a 2kb fragment downstream of *rocRp* (DeltaRocRlike_P3 and DeltaRocRlike_P4). Isolate HL 0640 3009 was naturally transformed with the PCR product and transformants were selected on kanamycin. Correct insertion of the kan-mazF cassette was verified by PCR. Second, the strain was naturally transformed with a PCR product creating a internal deletion of *rocRp*. The PCR product consists of the assembly of two PCR products corresponding to the upstream (primers DeltaRocRlike_P2d and DeltaRocRlike_P3d) and downstream sequence of *rocRp* (Primers DeltaRocRlike_P1 and DeltaRocRlike_P4). Transformants were selected on IPTG and were checked for sensitivity to kanamycin. Plasmid extraction and PCR analysis confirmed that pLPL was still present in the isolate and that *rocRp* was effectively deleted. (B) Conjugative transfer of pLPL^3009KF^ to JR32, Paris and other clinical isolates. Streptomycin-resistant mutant of isolates of LG-0712-2011, Paris and HL-0638-5028 were obtained by plating 1 mL of culture on CYE plated containing streptomycin and isolating spontaneous resistant mutants. JR32 is readily resistant to streptomycin. The streptomycin resistant recipients were inoculated in 3 mL of AYE medium together with the streptomycin-sensitive strain 3009 pLPL ^3009KF^ at OD=0.1 in 13-mL tubes. The cultures were grown for 48H at 30°C in a shaking incubator and 100 µL were then plated on CYE containing kanamycin and streptomycin. Cultures of recipients or donor alone did not give rise to colonies. Presence of the pLPL^3009KF^ plasmid in the transconjugants was confirmed by plasmid extraction and PCR analysis. The donor isolate, HL-0604-3009 shows an atypical growth curve with a stationary phase (OD=3.9-4.2) lower than most *L. pneumophila* isolates (OD=5.5-6.0), including the recipients. This phenotype was used to verify that kanamycin and streptomycin resistant mutants corresponded to transconjugants rather than spontaneous streptomycin resistant mutants of the donor. (C) Construction of Paris strain carrying *rocRp* on plasmid pLPP. The *rocRp* gene was introduced in the pseudogene *plpp0110* of pLPP by natural transformation of the Paris strain. A PCR product was assembled using 2 kb of sequence upstream of *plpp0110* (primers plpp0110_P1 and plpp0110_P2g), the *rocRp* gene (primers KP1 and KP2), the gentamicin resistance gene (primers gnt-F and gnt-R) and 2k of sequence downstream of *plpp0110* (primers plpp0110_P3g and plpp0110_P4). The assembled PCR product was added to a liquid culture of the Paris strain and incubated 24h at 30°C for natural transformation to occur. Transformants were selected on CYE plates with gentamicin. Presence of the *rocRp* gene in pLPP was confirmed by PCR. (D) Construction of strain Paris_H1. Paris_H1 carries the *sfgfp* gene, encoding superfolder GFP, under the strong promoter J23119 (iGEM part Bba_J23119) inserted in the pseudogene *lpp0858a*. The strain was obtained by natural transformation of a PCR product assembled from four PCR products: a PCR consisting of 2 kb upstream of the targeted insertion site (primers lpp0858_P1 and lpp0858_P2), a PCR product of the *sfgfp* gene under the stong promoter J23119 (primers J23119 and termR), a PCR product of the gentamicin resistance gene (primers gnt-F and Cassette_Gm_Rv) and a PCR product of 2 kb of sequence downstream of the targeted insertion site (primers lpp0858_P7 and lpp0858_P8).

### Transformation assay of clinical isolates and GWAS

Clinical isolates listed in Table S1 were tested for natural transformability in 96-well plates. From a culture on CYE plate, isolates were grow in 100 µL of AYE in 96-well plates sealed with oxygen-permeable membrane (Sigma-Aldrich) at 30°C with shaking at 200 rpm (Infors HT) for 24H. Cultures were homogenized by pipetting and 2 µL were transferred to 100 µL of AYE containing 200 ng/µL of PCR product (primers rpsL_Fw and rpsL_Rv) of the *rpsL* gene region of the Paris_S strain. The plates were sealed with an oxygen-permeable membrane and incubated at 30°C in a shaking Thermomixer (Eppendorf) at 600 rpm. Control experiments were run in parallel, in the same conditions but without added DNA. After 48h, 10 µL of cultures were then spotted on CYE plates containing streptomycin and incubated at 37°C for three days. Each 96-well plate contains a transformable control strain (Paris) and a control streptomycin-resistant strain (JR32). Transformation was scored as a function of the approximate number of colonies that developed in the spot (no colony, score 0; 1 to 9 colonies, score 1; 10-50 colonies, score 2; so many colonies that the spot appears smooth, score 3). Transformation scores were determined four times independently. For each experiment, a score was retained only if it was superior to the score determined in the no DNA condition. A median score was then calculated for the n number of determinations that met this criteria. Phylogenetic relationship were determined using core-genome MLST generated with chewBBACA (59) and vizualised using GrapeTree (60). GWAS were carried out using DBGWAS 0.5.2 (42) on a binary matrix of non-transformable (NT, score 0-0.5) and transformable (T, score 1-3) phenotypes. Lineage effect was tested by DBGWAS by providing a phylogenetic tree based on whole genome assemblies using parsnp (61).

### Quantitative transformation assays

Natural transformation assays were conducted on cultures in AYE. The tested strains were inoculated at OD=0.1-0.2 in 3 mL of AYE in 13 mL tubes. One microgram of transforming DNA was added and the cultures were incubated at 30°C with shaking for 24H. Different transforming DNA were used depending on the resistance markers carried by the tested strains. A PCR product of encompassing the *rpsL* gene from the Paris_S strain was used for streptomycin-sensitive strains. Alternatively, transformation was tested with a non-replicative circular DNA carrying the kanamycin resistance gene inserted in the non-essential *ihfB* gene (pGEM-ihfB::kan) (44). For strains resistant to both kanamycin and streptomycin (*i.e.*, Paris_S pLPP::*rocRp*) transformation was tested with genomic DNA of strain Paris_H1 which carries the gentamicin resistance gene inserted in the pseusdogene *lpp0858a*. Transformation with genomic DNA typically yields 10-to 100-fold lower transformation frequencies than PCR product or plasmid. Transformation frequencies were obtained by calculating the ratio of transformants CFU over total CFU counts, determined by plating serial dilution of the cultures on selective and non-selective CYE plates. All transformation assays were performed at least three times independently, several days or weeks apart.

### Gene expression analysis by northern blot

Total RNA from bacterial cultures was extracted according to a previously described procedure (38). Briefly, bacterial cultures (1 mL) were mixed to an equal volume of ice-cold methanol, pelleted and lysed in 50 µl of RNAsnap buffer (18 mM EDTA, 0.025% SDS, 95% formamide). RNA was then extracted using a tri-reagent solution (acid guanidinium thiocyanate-phenol-chloroform) and isopropanol-precipitated. One to two micrograms of total RNA in denaturing buffer were loaded per lane of a denaturing Tris/Borate/EDTA (TBE)-urea 8% acrylamide gel. Ethidium-bromide staining of ribosomal RNA was used to check for equal loading of the lanes. RNA was transferred to a nylon membrane and cross-linked by UV irradiation. Membranes were hybridized at 42 °C with 5 nM of a 5’-biotinylated oligonucleotide probe (Table S4) in ULTRAhyb Ultrasensitive Hybridization Buffer (Ambion) and then washed according to the manufacturer’s instructions. Membranes were developed using HRP-conjugated streptavidin and enhanced luminol substrate (Pierce). Luminescence signals were acquired using an imaging workstation equipped with a charge-coupled device camera (Vilber-Lourmat).

### Global gene expression profiling by RNAseq

Total RNA from bacterial cultures grown to OD600=2 at 30°C were extracted as described above. RNA samples were treated with DNAse I and purified on silica-based columns (Zymo Research). Following ribosomal RNA depletion (Ribo-Zero, Epicentre), strand-specific cDNA libraries were prepared and sequenced on an HiSeq platform (Illumina) with paired-end 150bp assay. Enriched transcripts were determined using DESeq2 (62).

### 5’/3’ RACE of RocR and RocRp

5’/3’ RACE was performed essentially as previously described (63). Total RNA (5 µg) was treated with RppH (New England Biolabs) for 1 h at 37°C. After purification using the DirectZol kit (ZymoResearch), RNA was circularized with T4 RNA ligase overnight at 16°C. The ligase was inactivated by a 15 min-incubation at 70°C and RNA was purified using the DirectZol kit. RocR and RocRp were then reverse-transcribed with a specific primer (LA124) using the RevertAid H Minus kit (ThermoFischer). The PrimeSTAR Max (TaKaRa) was used to PCR-amplify either RocR (primers LA124/LA125) and/or RocRp (primers LA124/126). The PCR products were cloned using the CloneJET system (ThermoFischer) and sequenced.

### Purification and relative quantification of RocC-bound RNAs

Bacterial cultures at the optical densities indicated in the text were fixed with 1% formaldehyde for 30 min. Pelleted bacterial cells were resuspended in lysis buffer (50 mM HEPES-KOH pH7.5, 150mM NaCl, 1 mM EDTA, 1% Triton X-100, 0.1% Na-deoxycholate) and sonicated at 4°C. Lysates were incubated with Dynabeads Protein A magnetic beads (Invitrogen) coated with rabbit-raised affinity-purified antibodies directed against RocC (38). Following washing steps, RNA were eluted from the magnetic beads by extraction with a tri-reagent solution (acid guanidinium thiocyanate-phenol-chloroform) and isopropanol-precipitated. RNA were reverse-transcribed (RevertAid, Thermo) in a single reaction with a mix of two primers specific for RocRp and RocR (primers RT19-rocRlike and RT20-rocR). Both primers carry a 5’ extension which was then used for the simultaneous PCR amplification of RocR and RocRp with primers RRl17F and RRl20R. The resulting PCR product, a mix of RocR (86 bp) and RocRp (85 bp), were purified and digested with NruI, which cleaves only RocRp, giving two fragments of 42 and 43 bp. The restriction digests were quantified by capillary electrophoresis (TapeStation, Agilent).

### mRNA decay and RNA half-life determination

Bacterial cultures at optical densities indicated in the text and figures were treated with 100 µg/ml of rifampicin to stop transcription. RNA were extracted as described above at the timepoints indicated in the figure legends. RocR and RocRp levels were detected by Northern-blot analysis and quantitated by densitometry with ImageJ. Values of RocR and RocRp levels were normalized to levels of rRNA. Values at t=0 were normalized to 1. Data were fit to a first-order exponential decay with the Qtiplot software.

### Genomes sequencing, assembly and analysis

Environmental isolates of *L. geestiana* HL-0438-4026 and *L. israelensis* HL-0427-4011 were sequenced using both Illumina and Oxford Nanopore technologies. For Illumina sequencing, whole genomes were sequenced in paired-end 2×300bp on a Miseq sequencer using Nextera XT kit according to manufacturer’s instructions. Whole genomes were sequenced using the Rapid barcoding kit on a MinION sequencer according to manufacturer’s instructions (Oxford Nanopore). Illumina reads were trimmed for low quality nucleotides and adapters removing using trimmomatic 0.36 (64). Nanopore reads were base-called and demultiplexed using guppy (Oxford Nanopore). Illumina and Nanopore reads were then used for short reads/long reads hybrid assembly using Unicycler v0.4.6 (65). Similarly, the genomes of *L. israelensis* ATCC43119 and *L. geestiana* ATCC49504 were assembled from raw Illumina and Pacbio sequencing data downloaded from NCBI SRA database. All genomes were annotated using prokka 1.13 (66). Presence of *rocR* and *rocRp* sequences in *Legionella* genus genomes were investigated using local blastn on all the 835 *Legionella*/*Fluoribacter*/*Tatlockia* genome assemblies (697 *Lp*, 138 non-*Lp*) available from the NCBI assembly database on March 6^th^, 2019. Detection of conjugative elements was performed using MacSyFinder (67) and the ConjScan module (46).

### Accession numbers

Genomes assemblies reported in this paper have been deposited to Genbank (Bioproject accession number PRJNA528641). RNA-Seq raw reads were deposited to the European Nucleotide Archive (study accession number PRJEB31835). Sequence data of all clinical isolates used in this study were previously published and are available from the European Nucleotide Archive under accession number PRJEB15241 (40).

## Acknowledgments

This work was supported in part by a grant from Agence Nationale de la Recherche to LA (Project RNAchap, ANR-17-CE11-0009-01). This work was performed within the framework of the LABEX ECOFECT (ANR-11-LABX-0048) of Université de Lyon, within the program “Investissements d’Avenir” (ANR-11-IDEX-0007) operated by the French National Research Agency (ANR). We are grateful to Magali Jaillard and Laurent Jacob for introducing us to the DBGWAS tool and for all subsequent discussions about its use in our project. We thank Louise Kindingstad (Stavanger University Hospital, Norway) for providing the *L. israelensis* isolate.

