## Supplemental Table and Figures for "Diverse conjugative elements silence natural transformation in *Legionella* species"

**Supporting information**

Isabelle Durieux[1,#], Christophe Ginevra[2,3,#], Laetitia Attaiech[1], Kevin Picq[1], Pierre-Alexandre Juan[1], Sophie Jarraud[2,3], Xavier Charpentier[1]*

[1] CIRI, Centre International de Recherche en Infectiologie, Team "Horizontal gene transfer in bacterial pathogens", Inserm, U1111, Université Claude Bernard Lyon 1, CNRS, UMR5308, École Normale Supérieure de Lyon, Univ Lyon, 69100, Villeurbanne, France
[2] CIRI, Centre International de Recherche en Infectiologie, Team "Pathogenesis of *Legionella*", Inserm, U1111, Université Claude Bernard Lyon 1, CNRS, UMR5308, École Normale Supérieure de Lyon, Univ Lyon, 69008, Villeurbanne, France
[3] Centre National de Référence des Légionelles, Centre de Biologie et de Pathologie Est, 59 Boulevard Pinel, 69677 Bron Cedex, France

### These authors contributed equally

**Table S1. Transformation scores of strains and clinical isolates for GWAS.**

Natural transformability was tested in 96 well plates. Transforming DNA (a 4kb PCR product encompassing the *rspL* gene of the streptomycin-resistant mutant Paris_S) is added to 100 µl of AYE inoculated with the isolate. The culture was incubated 48H at 30°C with orbital shaking. A 10 µl sample was then spotted on a CYE plate containing streptomycin (50 µg/mL) and incubated for 3 days at 37°C. Transformation was scored as a function of the approximate number of colonies that developed in the spot (score 0 = no colony; score 1 = 1 to 9 colonies; score 2 = more than 10 colonies but still distinguishable; score 3 = so many colonies that the spot appears smooth). Transformation scores was determined four times independently. A control without DNA is conducted in parallel. For each experiment, a score is retained only if it is superior to the score determined in the no DNA conditions. The median score is reported for the n number of determinations that met this criteria. Isolates are considered transformable (T) if the median score is higher than 0.5, otherwise they are considered non-transformable (NT). ST, presence of a pLPL plasmid and *rocRp*, and the average transformation frequencies determined using the ihfB::kan marker (shown in Fig. 1A) are also reported.

| Isolate | ST | pLPL | rocRp | MEDIAN SCORE | T/NT | n= | TF (ihfB::kan) |
|---|---|---|---|---|---|---|---|
| Paris | 1 | | | 2 | T | 4 | 1.2E-05 |
| HL 0618 5018 | 23 | | | 2 | T | 4 | |
| HL 0620 4017 | 40 | | | 0 | NT | 4 | |
| HL 0622 5007 | 23 | | | 1 | T | 3 | |
| HL 0622 5017 | 40 | | | 0 | NT | 4 | |
| HL 0623 2008 | 23 | | | 1.5 | T | 4 | |
| HL 0624 5009 | 23 | | | 2 | T | 4 | 1.3E-07 |
| HL 0626 3008 | 664 | | | 1 | T | 4 | 2.2E-07 |
| HL 0626 3031 | 317 | | | 2 | T | 4 | |
| HL 0627 4016 | 700 | | | 1 | T | 3 | |
| HL 0629 2024 | 44 | | | 1 | T | 4 | |
| HL 0629 4018 | 23 | | | 0 | NT | 3 | 5.8E-09 |
| HL 0631 1018 | 146 | | | 0.5 | NT | 4 | 3.5E-08 |
| HL 0631 2017 | 1 | | | 2.5 | T | 4 | |
| HL 0631 2018 | 700 | | | 0.5 | NT | 4 | |
| HL 0631 3013 | 455 | | | 0 | NT | 4 | 5.6E-08 |
| HL 0633 3016 | 26 | | | 0 | NT | 3 | |
| HL 0633 5004 | 23 | + | + | 0 | NT | 3 | |
| HL 0634 5015 | 441 | | | 2.5 | T | 4 | |
| HL 0634 5016 | 40 | | | 0 | NT | 3 | |
| HL 0635 1002 | 438 | | | 0 | NT | 4 | |
| HL 0635 2011 | 23 | | | 2 | T | 4 | |
| HL 0635 3025 | 47 | | | 0 | NT | 4 | |
| HL 0635 5023 | 62 | | | 2 | T | 4 | |
| HL 0636 2015 | 47 | | | 0 | NT | 4 | |
| HL 0636 4015 | 23 | | | 1.5 | T | 4 | 1.9E-07 |
| HL 0636 5016 | 1 | | | 2.5 | T | 4 | |
| HL 0637 3020 | 146 | | | 1.5 | T | 4 | |
| HL 0637 3028 | 65 | | | 0 | NT | 4 | |
| HL 0637 4022 | 47 | | | 0 | NT | 4 | |
| HL 0637 4023 | 146 | | | 1.5 | T | 4 | |

| | | | | | | | |
|---|---|---|---|---|---|---|---|
| HL 0637 4025 | 23 | + | - | 1.5 | T | 4 | |
| HL 0638 5028 | 1 | | | 2 | T | 4 | 1.6E-05 |
| HL 0638 7006 | 448 | | | 2 | T | 4 | |
| HL 0639 2009 | 1 | | | 2 | T | 4 | |
| HL 0639 2021 | 1 | | | 2.5 | T | 4 | |
| HL 0639 4022 | 624 | | | 0 | NT | 4 | |
| HL 0639 4025 | 20 | | | 0 | NT | 4 | |
| HL 0639 5028 | 47 | | | 0 | NT | 4 | |
| HL 0640 1010 | 47 | | | 0.5 | NT | 4 | |
| HL 0640 1051 | 47 | | | 0 | NT | 3 | |
| HL 0640 3004 | 47 | | | 0.5 | NT | 2 | |
| HL 0640 3009 | 23 | + | + | 0.5 | NT | 4 | 2.0E-09 |
| HL 0640 4012 | 146 | + | + | 0 | NT | 3 | <1E-9 |
| HL 0640 4016 | 94 | | | 0 | NT | 4 | 1.7E-08 |
| HL 0640 5008 | 62 | | | 2.5 | T | 4 | 4.8E-06 |
| HL 0641 2019 | 20 | | | 0 | NT | 2 | |
| HL 0641 3006 | 701 | + | + | 0.5 | NT | 4 | |
| HL 0641 3007 | 20 | | | 0 | NT | 4 | |
| HL 0641 5012 | 177 | | | 2 | T | 4 | |
| HL 0641 5021 | 9 | | | 0 | NT | 4 | 1.2E-08 |
| HL 0641 5022 | 207 | | | 2 | T | 4 | |
| HL 0642 1018 | 259 | | | 0 | NT | 4 | |
| HL 0642 5019 | 444 | | | 0.5 | NT | 4 | 1.3E-08 |
| HL 0642 5020 | 23 | | | 0 | NT | 4 | |
| HL 0642 5021 | 75 | | | 2 | T | 4 | |
| HL 0642 5022 | 47 | | | 0 | NT | 3 | |
| HL 0643 3031 | 146 | + | + | 0 | NT | 4 | 4.0E-09 |
| HL 0644 2011 | 1 | | | 3 | T | 3 | |
| HL 0645 2023 | 62 | | | 3 | T | 4 | |
| HL 0646 1021 | 65 | | | 0 | NT | 4 | |
| HL 0646 2023 | 37 | | | 0 | NT | 4 | |
| HL 0646 5011 | 23 | | | 2.5 | T | 4 | |
| HL 0647 2033 | 181 | | | 0 | NT | 4 | 6.9E-09 |
| HL 0647 5014 | 20 | + | + | 0 | NT | 3 | 3.4E-09 |
| HL 0648 2012 | 59 | | | 2 | T | 3 | |
| HL 0648 5020 | 9 | | | 0 | NT | 3 | |
| HL 0648 5023 | 42 | | | 2 | T | 4 | |
| HL 0650 4059 | 20 | | | 0 | NT | 4 | |
| HL 0651 1014 | 23 | | | 3 | T | 4 | |
| HL 0652 4035 | 96 | | | 1.5 | T | 4 | |
| HL 0701 3035 | 23 | | | 3 | T | 4 | |
| HL 0702 5017 | 1 | | | 2.5 | T | 4 | |

| | | | | | | | |
|---|---|---|---|---|---|---|---|
| HL 0703 3038 | 23 | | | 2 | T | 4 | |
| HL 0703 5020 | 23 | + | - | 2.5 | T | 4 | |
| HL 0705 5011 | 47 | | | 0 | NT | 4 | |
| HL 0707 2036 | 224 | | | 0 | NT | 4 | 6.2E-09 |
| HL 0709 3014 | 18 | | | 3 | T | 4 | 4.1E-06 |
| HL 0710 2013 | 75 | | | 2 | T | 4 | |
| HL 0710 5012 | 47 | | | 0 | NT | 4 | |
| LG 0712 2011 | 1 | | | 3 | T | 4 | 3.1E-05 |
| LG 0713 5012 | 1 | | | 3 | T | 4 | |
| LG 0714 2022 | 59 | | | 0 | NT | 3 | |
| LG 0714 4016 | 20 | | | 0 | NT | 4 | |
| LG 0715 5014 | 23 | | | 2 | T | 4 | |
| LG 0716 2016 | 1 | | | 3 | T | 4 | |
| LG 0717 5021 | 40 | | | 1 | T | 4 | |
| LG 0721 1006 | 702 | | | 1 | T | 4 | 1.8E-06 |
| LG 0721 5005 | 23 | | | 2 | T | 4 | |
| LG 0723 4019 | 23 | + | + | 0 | NT | 3 | |
| LG 0723 4026 | 23 | + | + | 0.5 | NT | 4 | |
| LG 0724 2038 | 23 | | | 2.5 | T | 4 | 4.5E-06 |
| LG 0724 2039 | 82 | + | + | 0 | NT | 3 | |
| LG 0724 3029 | 703 | + | + | 1 | T | 4 | |
| LG 0724 5003 | 6 | | | 2 | T | 4 | 1.5E-06 |
| LG 0724 5013 | 708 | | | 0 | NT | 4 | <1E-9 |
| LG 0725 2030 | 1 | | | 2 | T | 4 | |
| LG 0725 4014 | 1 | | | 3 | T | 4 | |
| LG 0725 5007 | 94 | | | 0 | NT | 4 | 2.7E-08 |
| LG 0726 2016 | 146 | | | 1 | T | 3 | |
| LG 0726 4009 | 94 | | | 0 | NT | 3 | |
| LG 0727 3031 | 96 | | | 0.5 | NT | 4 | |
| LG 0727 4020 | 37 | | | 1 | T | 4 | |
| LG 0727 5009 | 146 | | | 1.5 | T | 4 | |
| LG 0727 5020 | 23 | | | 2 | T | 4 | 7.8E-07 |
| LG 0727 5021 | 77 | | | 0 | NT | 3 | |
| LG 0728 1015 | 44 | | | 1.5 | T | 4 | |
| LG 0728 3011 | 23 | + | + | 0 | NT | 1 | |
| LG 0729 4019 | 704 | | | 2 | T | 4 | |
| LG 0751 3022 | 47 | | | 0 | NT | 3 | |
| OLDA | ND | | | 2 | T | 4 | |
| 130B | ND | | | 1.5 | T | 4 | |
| Lens | 15 | + | + | 0 | NT | 4 | <1E-9 |

**Table S2. RNAseq analysis of the 3009ΔpLPL and 3009ΔrocRp strains relative to the wild-type 3009 isolate (HL 0640 3009).**

Total RNA from three independent bacterial cultures grown to an OD 600 of 1.8-2.0 at 30°C was extracted, purified, and sequenced (Illumina). The number of reads was normalized, and the statistical significance ($P_{adj}$) of the fold change (FC) was determined with DEseq2. List is limited to chromosomal genes for which $log_2$FC is >1 in either of the two comparisons.

| Gene Identifiers | log2FC ΔpLPL vs WT | log2FC ΔrocRp vs WT | $P_{adj}$ ΔpLPL vs WT | $P_{adj}$ ΔrocRp vs WT | gene names | lpp number in the Paris strain |
|---|---|---|---|---|---|---|
| STR10_v2_170063 | 3.32 | 2.00 | 3.69E-99 | 4.66E-34 | comM | lpp0640 |
| STR10_v2_50169 | 3.84 | 2.47 | 2.40E-80 | 8.76E-32 | comEA | lpp0872 |
| STR10_v2_170020 | 2.72 | 1.75 | 4.28E-44 | 3.97E-17 | comEC | lpp0680 |
| STR10_v2_270143 | 2.93 | 2.17 | 5.23E-41 | 6.50E-21 | | lpp2554 |
| STR10_v2_270142 | 2.01 | 1.32 | 4.00E-26 | 3.32E-10 | radC | lpp2553 |
| STR10_v2_110039 | 1.65 | 1.32 | 1.53E-13 | 1.67E-07 | | lpp1977 |
| STR10_v2_160037 | 1.47 | 1.02 | 1.31E-09 | 1.52E-03 | comF | lpp2280 |
| STR10_v2_110037 | 1.08 | 0.49 | 3.26E-06 | 2.44E-01 | | lpp1976 |
| STR10_v2_50147 | 1.23 | 0.95 | 8.86E-06 | 1.68E-02 | pilE | lpp0851 |

**Table S3. Bacterial strains used or generated in this study.**
A comprehensive list of *L. pneumophil*a clinical isolates used in this study are listed in Table S1. Abbreviations of antibioresistance are as follows: Gent for Gentamycin, Kan for Kanamycin and Strep for Streptomycin.

| **Strains** | **Genotype and/or description** | **Reference** |
|---|---|---|
| Paris WT | Paris outbreak isolate CIP107629 | CNR Lyon |
| Paris_S | Spontaneous streptomycin-resistant mutant of Paris; $Strep^R$ | This study |
| Paris H1 | *lpp0858a*::(J23119-sfGFP-*aacC*1); $Gent^R$ | This study |
| HL-0640-3009 | Clinical isolate | CNR Lyon |
| 3009 $pLPL^{3009KF}$ | pLPL::(*lacI*q, Ptac-*mazF*, *npt*II); $Kan^R$, $IPTG^S$ | This study |
| 3009ΔpLPL | 3009 $pLPL^{3009KF}$ that lost pLPL; $Kan^S$, $IPTG^R$ | This study |
| 3009 ΔrocRp::kan-mazF | HL_0640_3009; ΔrocRp::(*lacI*q, Ptac-*mazF*, *npt*II); $Kan^R$, $IPTG^S$ | This study |
| 3009Δ*rocRp* | HL_0640_3009; ΔrocRp; $Kan^S$, $IPTG^R$ | This study |
| Paris_S $pLPL^{3009KF}$ | Transconjugant of 3009 $pLPL^{3009KF}$ and Paris_S; $Strep^R$, $Kan^R$, $IPTG^S$ | This study |
| Paris $rocC_{TAA}$ | Paris; $rocC_{TAA}$ (=lpp0148); *rocC* allele with a premature stop codon TAA | Juan et al., 2015 |
| Paris Δ*rocR* | Paris; Δ*rocR* | Attaiech et al., 2016 |
| Paris pLPP::*rocRp* | Paris; pLPP::*(rocRp-aacC*1); $Gent^R$ | This study |
| $rocC_{TAA}$ pLPP::*rocRp* | $rocC_{TAA}$ pLPP::*(rocRp-aacC*1); $Gent^R$ | This study |
| Δ*rocR* pLPP::*rocRp* | Δ*rocR* pLPP::*(rocRp-aacC*1); $Gent^R$ | This study |
| LG-0712-2011_S | Spontaneous streptomycin-resistant mutant of LG-0712-2011; $Strep^R$ | This study |
| HL-0638-5028_S | Spontaneous streptomycin-resistant mutant of HL-0638-5028; $Strep^R$ | This study |
| LG-0712-2011_S $pLPL^{3009KF}$ | Transconjugant of 3009 $pLPL^{3009KF}$ and LG-0712-2011_S; $Strep^R$, $Kan^R$, $IPTG^S$ | This study |
| HL-0638-5028_S $pLPL^{3009KF}$ | Transconjugant of 3009 $pLPL^{3009KF}$ and HL-0638-5028_S; $Strep^R$, $Kan^R$, $IPTG^S$ | This study |
| HL-0438-2026 | Environmental isolate of *Legionella geestiana* | CNR Lyon |
| HL-0427-4011 | Environmental isolate of *Legionella israelensis* | CNR Lyon |
| 019060433301 | Environmental isolate of *Legionella israelensis* | Louise Kindingstad |
| JR32 | Philadelphia-1 derivative; $Strep^R$; r − m + | Sadosky et al., 1993 |

**Table S4. Oligonucleotides used in this study.**

| Oligonucleotide name | Sequence (5' to 3') | Used for |
|---|---|---|
| MazFk7-F | CGACTCACTATAGGGCGAATTGGGCCGCT TTCCAGTCGGGAAACCTG | PCR amplication of kan-mazF cassette |
| MazFk7-R | CATATGCCACCGACCCGAGCAAACCCGAA GAAGTTGTCCATATTGGCCAC | PCR amplication of kan-mazF cassette |
| DeltaRocRlike_P1 | CTGCAGCTGAATAAGCTGGGTTATCC | Inserting the kan-mazF cassette in rocRp gene |
| DeltaRocRlike_P2 | GGCCCAATTCGCCCTATAGTGAGTCGGTC CAACCTCGCTTGTTAAGCAAG | Inserting the kan-mazF cassette in rocRp gene |
| DeltaRocRlike_P3 | GGGTTTGCTCGGGTCGGTGGCATATGCCA GCCCTTTCTACTTTGAATAATCTAC | Inserting the kan-mazF cassette in rocRp gene |
| DeltaRocRlike_P4 | ATCATAATGAGGTGCTCACAAACAGTG | Inserting the kan-mazF cassette in rocRp gene |
| DeltaRocRlike_P2d | GTCCAACCTCGCTTGTTAAGCAAG | Deleting rocRp by removing the previously inserted kan-mazF cassette |
| DeltaRocRlike_P3d | CTTGCTTAACAAGCGAGGTTGGACCCAGC CCTTTCTACTTTGAATAATCTAC | Deleting rocRp by removing the previously inserted kan-mazF cassette |
| gnt-F | TTGCCCATGGACGCACACCGTG | PCR amplication of gentamicin cassette |
| gnt-R | CTCCCCGCGCGTTGGCCGATTCATTAAGTG CCACCTGGCGGCGTTG | PCR amplication of gentamicin cassette |
| KP1 | CTGACCCGGGTGAGCCTTACAATAAAGTT GG | PCR amplification of rocRp |
| KP2 | AATAGCGGCCGCAGCGAATAGGAGGATA GATG | PCR amplification of rocRp |
| plpp0110_P1 | GCCAAACGCTATCAGGAGTTACCTC | Inserting rocRp-gentR in gene plpp0110 of pLPP |
| plpp0110_P2g | GTTTCCACGGTGTGCGTCCATGGGCAAGA TTGTCCCATCTCCTTGAACC | Inserting rocRp-gentR in gene plpp0110 of pLPP |
| plpp0110_P3g | TAATGAATCGGCCAACGCGCGGGGAGCGA AGAATCAAGAATAACGCCAC | Inserting rocRp-gentR in gene plpp0110 of pLPP |
| plpp0110_P4 | CTGATGAGGCGCTGCGAAGAG | Inserting rocRp-gentR in gene plpp0110 of pLPP |
| plpp0110_P2_rocRlike | CATCTATCCTCCTATTCGCTGCGGCCGCTA TTGATTGTCCCATCTCCTTGAACC | Inserting rocRp-gentR in gene plpp0110 of pLPP |
| gnt-F_rocRlike | CCAACTTTATTGTAAGGCTCACCCGGGTCA GTTGCCCATGGACGCACACCGTG | Inserting rocRp-gentR in gene plpp0110 of pLPP |
| lpp0858_P1 | ATCCTTCTCATTTAGAGAGGGATGCT | Inserting P23119-sfGFP in lpp0858 |
| lpp0858_P2 | TAAGTTGGACTTTCCCTTACTGGCTTCGGA TGATTCCTGCAATGCT | Inserting P23119-sfGFP in lpp0858 |
| J23119_F | GCCAGTAAGGGAAAGTCCAACTTATTGAC AGCTAGCTCAGTCCTAGGTATAATTTCACA CAGGAAACAGAATTC | Inserting P23119-sfGFP in lpp0858 |

| | | |
|---|---|---|
| term_R | GTTTCCACGGTGTGCGTCCATGGGCAAACGCAAAAAGGCCATCCGTC | Inserting P23119-sfGFP in lpp0858 |
| gnt-F | TTGCCCATGGACGCACACCGTG | Inserting P23119-sfGFP in lpp0858 |
| Cassette_Gm_Rv | TTCCCCGAAAAGTGCCACCT | Inserting P23119-sfGFP in lpp0858 |
| lpp0858_P7 | AGGTGGCACTTTTCGGGGAAGTACAACGACGAGAGACCGCA | Inserting P23119-sfGFP in lpp0858 |
| lpp0858_P9 | TGTGCAGACATACCCTCCCATA | Inserting P23119-sfGFP in lpp0858 |
| rpsL_Fw | GCAGCTCCAGATGGCTCAATC | PCR amplication of rpsL region (4kb) |
| rpsL_Rv | CAACCATACATGTCCATATTGACCAC | PCR amplication of rpsL region (4kb) |
| mazF-pLPL_P1F | GGTATTCAGTTGATCAATTCCCTTCG | Inserting kan-mazF cassette in a pseudogene of pLPL[3009] |
| mazF-pLPL_P2F | GGCCCAATTCGCCCTATAGTGAGTCGACATCTGAGACTGATGAGTAACC | Inserting kan-mazF cassette in a pseudogene of pLPL[3009] |
| mazF-pLPL_P3F | GGGTTTGCTCGGGTCGGTGGCATATGCTGATCATCGATGGAACAGACAATTTTG | Inserting kan-mazF cassette in a pseudogene of pLPL[3009] |
| mazF-pLPL_P4F | GATGGTAGATACTTTGCATCTGGC | Inserting kan-mazF cassette in a pseudogene of pLPL[3009] |
| RT19-rocRlike | AGTGGGTCACAGCTTGATGAGAAAGGGCTGGTCGTGTG | RT-PCR RocC-bound RNA |
| RT20-rocR | AGTGGGTCACAGCTTGATGAGAAAGGGCCAATCAGTGTG | RT-PCR RocC-bound RNA |
| RRl17-F | GCGAGGTTGGACTTGCT | RT-PCR RocC-bound RNA |
| RRl20-R | AGTGGGTCACAGCTTGATG | RT-PCR RocC-bound RNA |
| LA124_RACE_RocR-R | TGACACATAGCAAGTCCAAC | RACE mapping |
| LA125_RACE_RocR-F | TTCCACTGGGTCAATTGGCG | RACE mapping |
| LA126_RACE_RocRlike-F | ATCCAATGGGTGTGGTCGCG | RACE mapping |
| comEA-NB2 | Biotin-CTACAAAACGCTGCCCTATACCCTTGACTTCCGCAAGCTCTTCCAGAG-Biotin | Northern-blot probe for dectection of comEA |
| LA59_rocR-small-NB | Biotin-TGTGTCGCCAATTGACCCAGTGGAATGACACATAGCAAGTCCAACCTC-Biotin | Northern-blot probe for dectection of RocR |
| rocR-pLPL-NB2 | Biotin-GCTGGTCGTGTGTCGCGACCACACCCATTGGATTGACACAA-Biotin | Northern-blot probe for dectection of RocRp |

**Table S5. Plasmids used in this study.**

| Plasmid name | Reference | Used for |
|---|---|---|
| pXDC18 | Charpentier et al., J. Bact. 2008. | Template for PCR of gentamicin resistance gene |
| pASG-1 | Godeux et al., J. Bact. 2019. | Template for PCR of sfgfp gene |
| pGEM-kan-mazF (pGEM-MK) | Bailo et al., Meth. Mol. Biol. 2019; Attaiech et al., PNAS. 2016 | Template for kan-mazF cassette |
| pGEM-ihfB::kan | Juan et al., Sci Rep, 2015; Attaiech et al., PNAS. 2016 | Natural transformation assay. Plasmid carrying a kanamycin resistance gene in a 4kb *L. pneumophila* chromosomal region encompassing the *ihfB* gene |

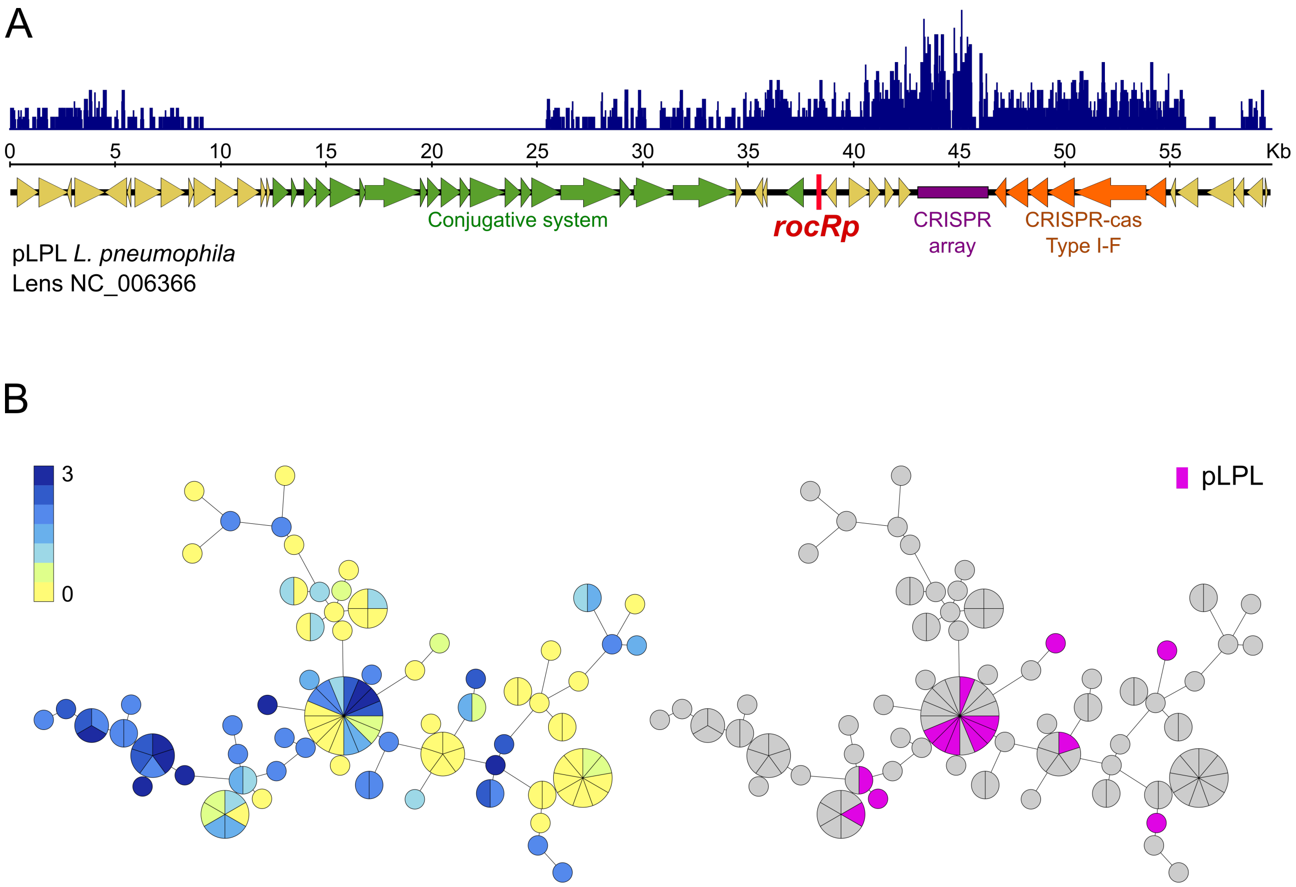


**Figure S1.** Plasmid pLPL associates with low levels of natural transformation. A. Linear map pf plasmid pLPL from the Lens isolate. Sequences of the 827 unambiguous sequences identified by DBGWAS as associating with the non-transformable phenotype (q<0.1) were mapped on the pLPL sequence (blue bars). B. Left panel: Semi-quantitative analysis of natural transformability of 113 isolates of *L. pneumophila*. Transformability was determined in 96-well format using a PCR product encompassing a *rpsL* allele conferring resistance to streptomycin. Genetic relationship were determined by cgMLST and visualized using a minimum spanning tree displaying transformation scores color coded from 0 (yellow) to 3 (dark blue). Right panel: Presence of the pLPL plasmid (pink) in the panel of 113 isolates tested for transformability.

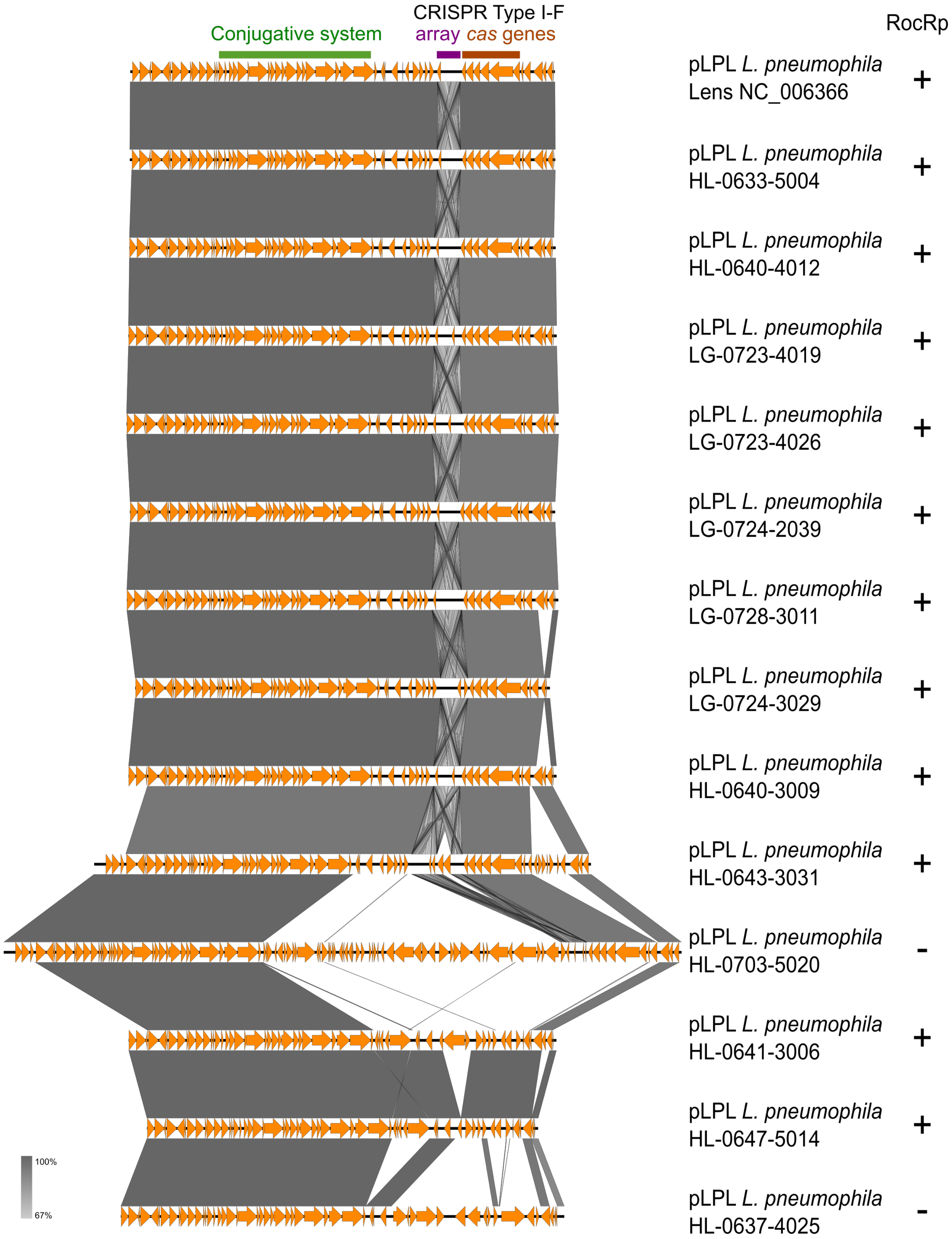


**Figure S2.** Multiple alignment of the 14 pLPL variants found in the panel of 113 clinical isolates investigated for their transformability levels. Plasmids pLPL from HL-0703-5020 and HL-0637-4025 show large indels and the absence of *rocRp*, which is consistent with their transformable phenotype.

Restriction digests on plasmids carried by mutants of the clinical isolate HL-0640-3009 (3009)

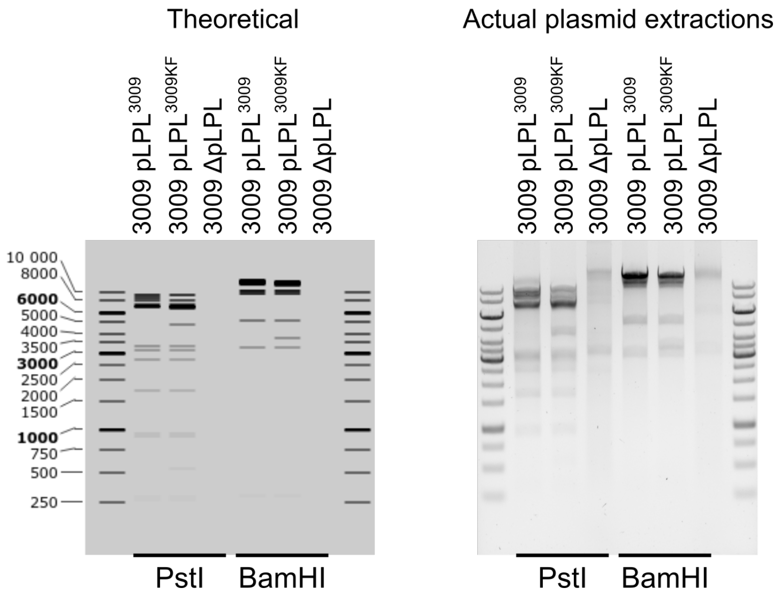


**Figure S3.** Restriction profiles of plasmid extraction from the clinical isolate HL-0640-3009 and mutants.
As expected, introduction of the kan-mazF cassette generates a different restriction profile due to the presence in the cassette of PstI and BamHI restriction sites. Plasmid pLPL is effectively absent from isolate 3009ΔpLPL. A smear is still visible due to the presence of contaminating genomic DNA in the plasmid preparation.

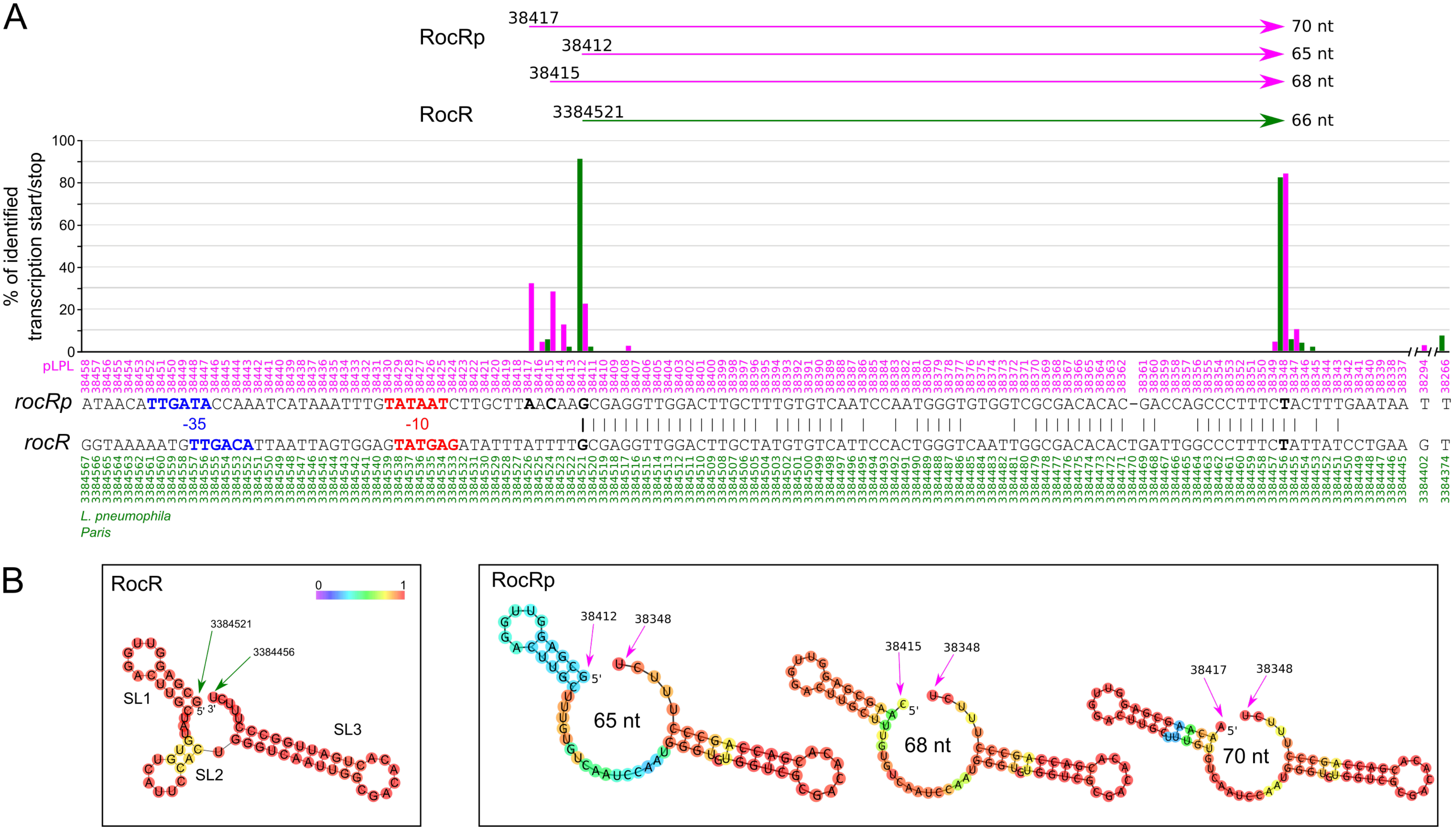


**Figure S4.** RACE determination of the 5' and 3' ends of RocR and RocRp. A. Relative abundance of identified 5' and 3'end of RocR (green) and RocRp (magenta). The sequence of the pLPL and the chromosome of the Paris strains are aligned based on the homology of the *rocR* and *rocRp* genes. Sequences highlighted in blue and green represent putative -10 and -35 box of the promoter, respectively. Nucleotides in bold font correspond to the mapped transcription start and termination sites. B. Secondary structures predicted by RNAfold of the RNA sequences identified for RocR and RocRp.

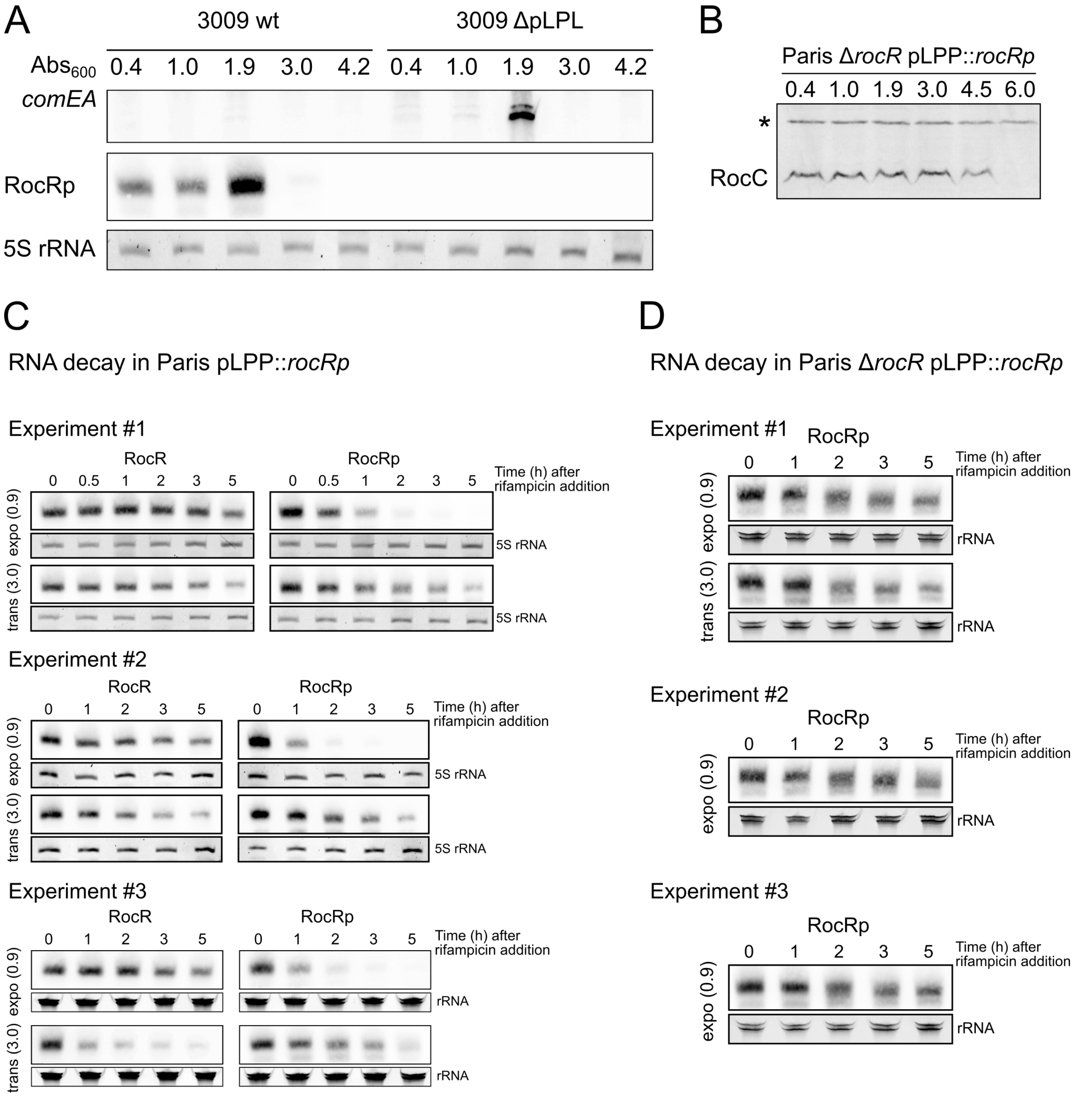


**Figure S5**. A. Northern-blot analysis of *comEA* and RocRp expression during growth of strains 3009 and 3009ΔpLPL in AYE medium at 30°C. Total RNA were extracted at the indicated optical densities (measured at 600 nm, $OD_{600}$) of the culture, and corresponding to the exponential growth phase (0.4 to 1.0), the transition phase (1.9) and early (3.0) and late stationary phase (4.2). Total RNA were separated on a 8% denaturing polyacrylamide gel. RocR and *comEA* transcripts were revealed with a biotinylated oligonucleotide probe. 5S rRNA was used as loading control. B. Western-blot analysis of RocC expression during growth at 30°C of the Paris strain carrying the *rocRp* gene on pLPP (Paris pLPP::*rocRp*). Total soluble proteins were extracted at the indicated optical densities ($OD_{600}$) of the culture, and corresponding to the exponential growth phase (0.4 to 1.0), the transition phase (1.9 and 3.0) and early (4.5) and late stationary phase (6.0). The star (*) indicates a cross-reactive band. C. Northern-blot experiments used to determine the half-life of RocR and RocRp in exponential phase (0.9) and transition phase (3.0) in the strain Paris pLPP::*rocRp*. When the cultures reached the indicated optical densities (0.9 and 3.0), rifampicin was added at 100µg/mL and samples were removed at the indicated times after addition of rifampicin. Total RNA was extracted and expression of RocR and RocRp was analysed by northern-blot. Signal intensities were determined using densitometry, normalized to rRNA levels and were fit to a first-order exponential decay with the Qtiplot software. Each experiment shown (#1, #2 and #3) were conducted on separate occasions, several days or weeks apart. D. Same as in C but with the strain Paris pLPP::*rocRp* deleted of the *rocR* gene.
